# Spatial Integration of Multi-Omics Data using the novel Multi-Omics Imaging Integration Toolset

**DOI:** 10.1101/2024.06.11.598306

**Authors:** Maximillian Wess, Maria K. Andersen, Elise Midtbust, Juan Carlos Cabellos Guillem, Trond Viset, Øystein Størkersen, Sebastian Krossa, Morten Beck Rye, May-Britt Tessem

## Abstract

To truly understand the cancer biology of heterogenous tumors in the context of precision medicine, it is crucial to use analytical methodology capable of capturing the complexities of multiple omics levels, as well as the spatial heterogeneity of cancer tissue. Different molecular imaging techniques, such as mass spectrometry imaging (MSI) and spatial transcriptomics (ST) achieve this goal by spatially detecting metabolites and mRNA, respectively. To take full analytical advantage of such multi-omics data, the individual measurements need to be integrated into one dataset. We present MIIT (Multi-Omics Imaging Integration Toolset), a Python framework for integrating spatially resolved multi-omics data. MIIT’s integration workflow consists of performing a grid projection of spatial omics data, registration of stained serial sections, and mapping of MSI-pixels to the spot resolution of Visium 10x ST data. For the registration of serial sections, we designed GreedyFHist, a registration algorithm based on the Greedy registration tool. We validated GreedyFHist on a dataset of 245 pairs of serial sections and reported an improved registration performance compared to a similar registration algorithm. As a proof of concept, we used MIIT to integrate ST and MSI data on cancer-free tissue from 7 prostate cancer patients and assessed the spot-wise correlation of a gene signature activity for citrate-spermine secretion derived from ST with citrate, spermine, and zinc levels obtained by MSI. We confirmed a significant correlation between gene signature activity and all three metabolites. To conclude, we developed a highly accurate, customizable, computational framework for integrating spatial omics technologies and for registration of serial tissue sections.

## Introduction

Detecting novel tissue biomarkers in cancer research is important for improving diagnosis and subsequent treatment choices. Spatial multi-omics has emerged as a promising methodological framework for identifying biomarkers within heterogenous cancer tissue. Whereas the traditional use of a single bulk omics technology, such as genomics, transcriptomics, metabolomics, or proteomics, could only capture one molecular level, multi-omics can elucidate complex biological processes occurring in the tumor tissue. Additionally, spatial omics technologies allow researchers to detect molecules within the spatial context of the tissue sample. In cancer research, spatially resolved methods are particularly powerful for analyzing the tumor microenvironment in situ containing a heterogeneous mixture of different cell types such as stroma cells, epithelial cells, and cancer cells of different aggressiveness. Spatial analysis are thus facilitating the identification of spatially defined molecular signatures,biomarkers and microenvironmental cellular interactions that otherwise would be lost if using traditional bulk molecular analysis. Two prominent examples for such spatial omics techniques are 10x Genomics’ Visium Spatial Gene Expression assay^1^ (ST) and mass spectrometry imaging (MSI) to detect spatial transcriptomics and spatial metabolomics data, respectively, together with tissue morphology. Even though this allows analyzing the link between single-spatial-omics and tissue morphology, to harness the full potential of such data and truly move towards spatial multi-omics, a reliable data integration methodology that accurately registers several spatial omics layers is required.

Collecting several different spatial omics data from the same tissue is often not possible due to incompatible sample preparation or destructive analyte extraction procedures, making it necessary to obtain serial sections, one for each omics layer, from the tissue of interest. Since standard histology staining can often be performed in the exact same section used for spatial-omics analysis, stained images can be used to find a registration between serial sections which are then applied to different spatial-omics modalities. We refer to registration as the process of transforming one image to the same coordinate system as another. Another challenge is that different spatial-omics platforms operate with different sampling resolution and organization. For instance, ST is organized as a sparse collection of equidistant spots of 50 µm diameter arranged in a hexagonal grid (*ST-spots*) and MSI consists of a dense map of pixels of 1 – 200 µm resolution (*MSI-pixels*). It is therefore necessary to establish a way of transforming between MSI-pixels and ST-spots after registration of stained images. ST and MSI have been integrated before, however these approaches have been performed semi-manual^2,3^, relied on proprietary software^3,4^. One alternative method performed ST and MSI on the same tissue section, removing the need for registration of serial sections. However, this protocol limits the number of omics modalities that can be used. Therefore, we need novel highly accurate end-to-end pipelines that can handle the registration of serial sections, can include several omics methodologies and can be automatic and openly accessible for the research community.

A significant challenge of registering serial sections is that the tissue composition between sections vary and potential deformation artifacts from sectioning can occur. This complicates an accurate registration between each neighboring section with increasing complexity as the distance between sections increases. Although several algorithms have been proposed to solve the task of registering serial sections^5–17^, they are largely based on FFPE tissue sectioned at close distance (e.g., 4-6 µm). FFPE sections are far less fragile and prone to artifacts compared to frozen tissue required for many spatial omics measurements, therefore requiring robust registration methods.

Our group has generated a comprehensive and complex spatial multi-omics dataset (“‘Tissue is the issue’: a multi-omics approach to improve prostate cancer diagnosis” (ERC: 758306)) as presented in Figure 1^18–20^. In this dataset ST and MSI were performed on serial sections from cylindrical tissue samples that have an axial sectioning distance of up to 100 µm, which increases the difficulty of a viable registration due to stronger effects of tissue heterogeneity and tissue damage. In this study, we have tackled the challenges of spatial multi-omics data integration by developing two open-source applications. With our tool GreedyFHist we could register stained heterogeneous serial sections that were up to 100 µm apart. GreedyFHist is compatible with most common image formats as well as ome.tiff and geojson^21^, and integrates well with QuPath^22^, a popular open-source software for digital pathology. Building further on this framework we developed Multi-Omics Imaging Integration Toolset (*MIIT*), an application capable of coordinating the different spatial resolutions and sampling organizations between different spatial modalities to create one spatial omics dataset. We used MIIT to define an end-to-end workflow for integrating ST and MSI by finding a robust registration between serial histology sections. Our integration works independently of the molecular properties measured, and new data types can be implemented to extend to other spatial omics technologies. Moreover, MIIT implements several utility functions for processing various spatial omics data types. To demonstrate the potential of MIIT, we investigated the correlation between a gene signature score for citrate and spermine secretion^23^ from ST data with metabolic measurements from MSI in glands of cancer-free tissue samples from prostate cancer patients. To calculate the gene signature scores we used the single sample Gene Set Enrichment Analysis (ssGSEA)^24^.

**Figure 1.**
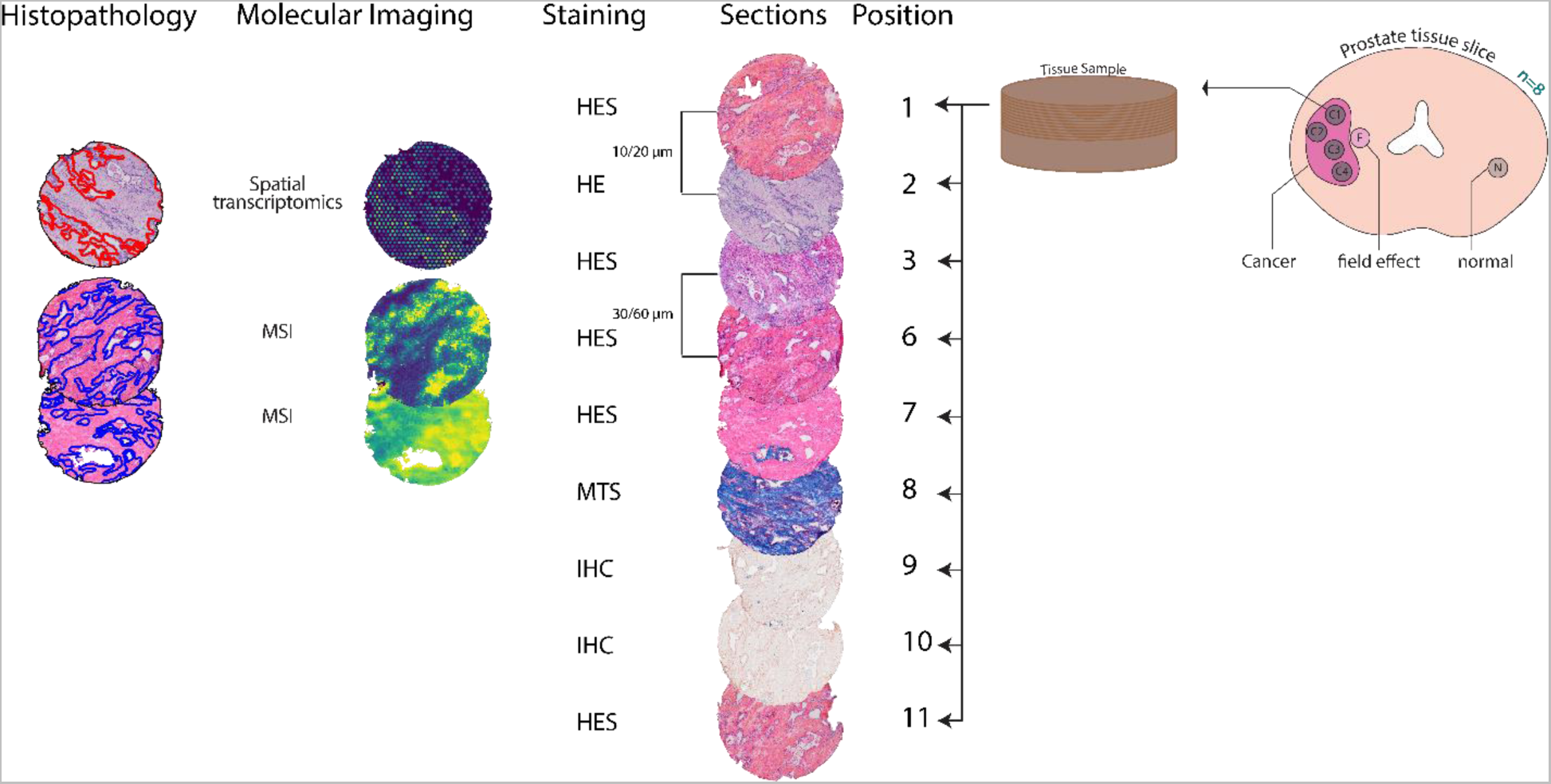
Overview of the ProstOmics spatial multi-omics dataset on prostate tissues. Sections of each core are numbered and processed in the same order. The sections used in this study are at positions 1-3 and 6-11 with a distance between two adjacent sections of 10 – 20 µm. The following staining techniques are used: Hematoxylin, Erythrosine and Saffron (*HES*); Hematoxylin and Eosin (*HE*); Mason Trichrome Staining (*MTS*); Immunohistochemistry (*IHC*). Spatial transcriptomics is applied to section 2, MSI in positive ion mode on section 6, and MSI in negative ion mode on section 7. Histopathology was evaluated for sections 2, 6, and 7.

## Results

Here we present a summary of GreedyFHist, our algorithm for registration of stained serial images and MIIT, the framework for integration of spatial-omics data. Detailed descriptions for both methods can be found in the methodology section.

### GreedyFHist for registration of histology images

GreedyFHist is our algorithm for registration of stained fresh frozen serial images. Image registration is the process of finding a transformation from one image (*moving image*) to another (*fixed image*) such that the transformation applied on the moving image results in a warped image which is aligned to the fixed image. *The main steps of the GreedyFHist algorithm are presented below and in Figure 2*.

**Figure 2.**
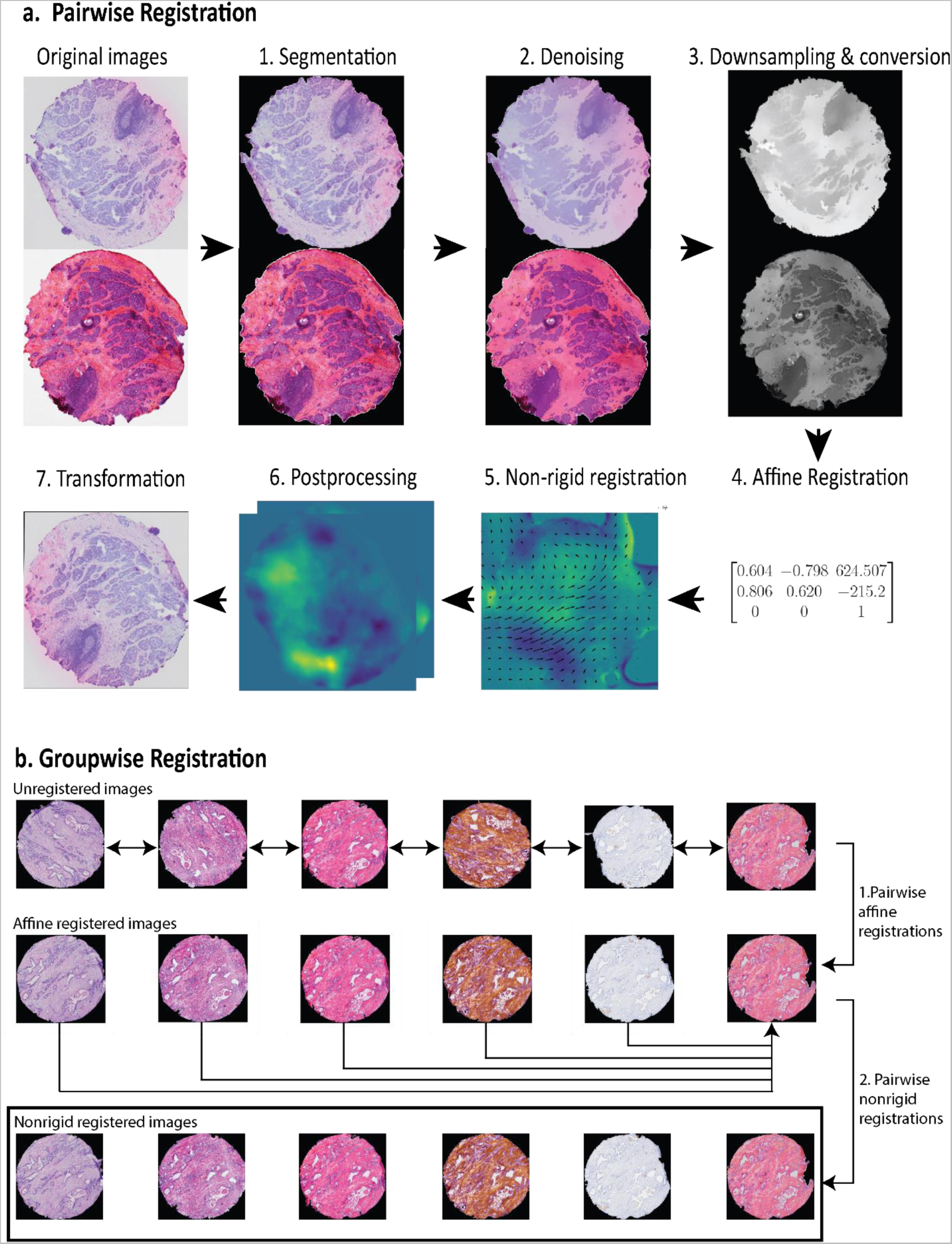
Overview of GreedyFHist’s registration. a) Registration between two images. In step 1, images are segmented from background to focus on tissue region. Then denoising (step 2) is applied to remove background while retaining major histological features. Grayscale conversion and downscaling (step 3) are used on the images to improve registration time. Next, affine (step 4) and non-rigid registration (step 5) are applied on preprocessed images through Greedy to compute transformation matrices. Transformation matrices are then rescaled to the original image’s resolution, composited into one transformation matrix (step 6) and applied to the moving image (step 7). b) **Groupwise registration.** To register a group of ordered images, the pairwise registration is applied on each consecutive pair of images. (step 1) Then a nonrigid registration is applied between each affine registered image and the fixed image (step 2).

Pairwise registration:

1. **Segmentation:** Images are first segmented to remove any background noise and extract the tissue area using a segmentation algorithm based on the YOLO8^25^ model.
2. **Denoising:** Then images are denoised to remove unnecessary image features while keeping major histological morphology features intact.
3. **Grayscale conversion & downscaling:** In the last preprocessing step, images are converted to grayscale color space and downscaled to 1024 x 1024 pixel resolution to reduce runtime of the registration.
4. **Affine registration**: During the affine registration, a global registration between moving and fixed image is computed using the diffeomorphic registration tool *Greedy*^26^.
5. **Nonrigid registration**: After the affine registration, Greedy is used to perform a nonrigid registration to align locally matching image features.
6. **Postprocessing**: The computed transformation matrices are rescaled to the moving and fixed image’s size and composited into one transformation matrix.
7. **Transformation**: Transformation matrices are used to transform images or pointset data from the moving image space to the fixed image space.

Groupwise registration:

In this mode, an ordered series of stained images is registered to a common fixed image. We denote the last image in the series as the fixed image.

1. **Pairwise affine registration:** On each pair of neighboring stained images a pairwise affine registration is performed and consecutively applied on each stained image, resulting in an affine registration for the whole image series.
2. **Nonrigid registration:** Then, a non-rigid registration between each image in the series and the fixed image is performed. As with the pairwise registration, each transformation is composited to reduce interpolation errors.

*Features:*

GreedyFHist supports the most common image formats including ome.tiff^27^ and has support for applying registration to spatial data in image, pointset and geojson^21^ format by which it interfaces with image analysis software such as QuPath. Because the task of registration for neighboring tissue sections can be applied in a variety of topics outside of the scope of this work, we provide GreedyFHist as a standalone application.

### MIIT (Multi-omics Imaging Integration Toolset)

MIIT (Multi-omics Imaging Integration Toolset) is a framework for integrating molecular imaging data. An illustration of MIIT’s workflow for the integration of ST and MSI can be found in Figure 3. By integrating we refer to the registration of sections and accurate transformation of resolution and granularity between different types of spatial omics data. In the following text, we use the term *section* to describe stained images, spatial omics data and histopathology annotations that correspond to each other. We explain MIIT’s integration workflow by highlighting how a s*ource section* analyzed with MSI is integrated to a *target section* analyzed with ST.

1. **Preprocessing**: Spatial omics data is registered with stained images and a grid projection of the molecular imaging data onto the image space of the stained image which allows accurate and efficient image transformations during registration.
2. **Registration**: Using the stained images as reference images the source section is registered to the image space of the target section using GreedyFHist.
3. **Transformation**: Spatial omics data from the source section is transformed to match the spatial organization of the target section’s molecular imaging data and, optionally, additional histology annotations.
4. **Export**: Integrated molecular imaging data from the source section is converted and exported to appropriate data formats for further analysis.

**Figure 3.**
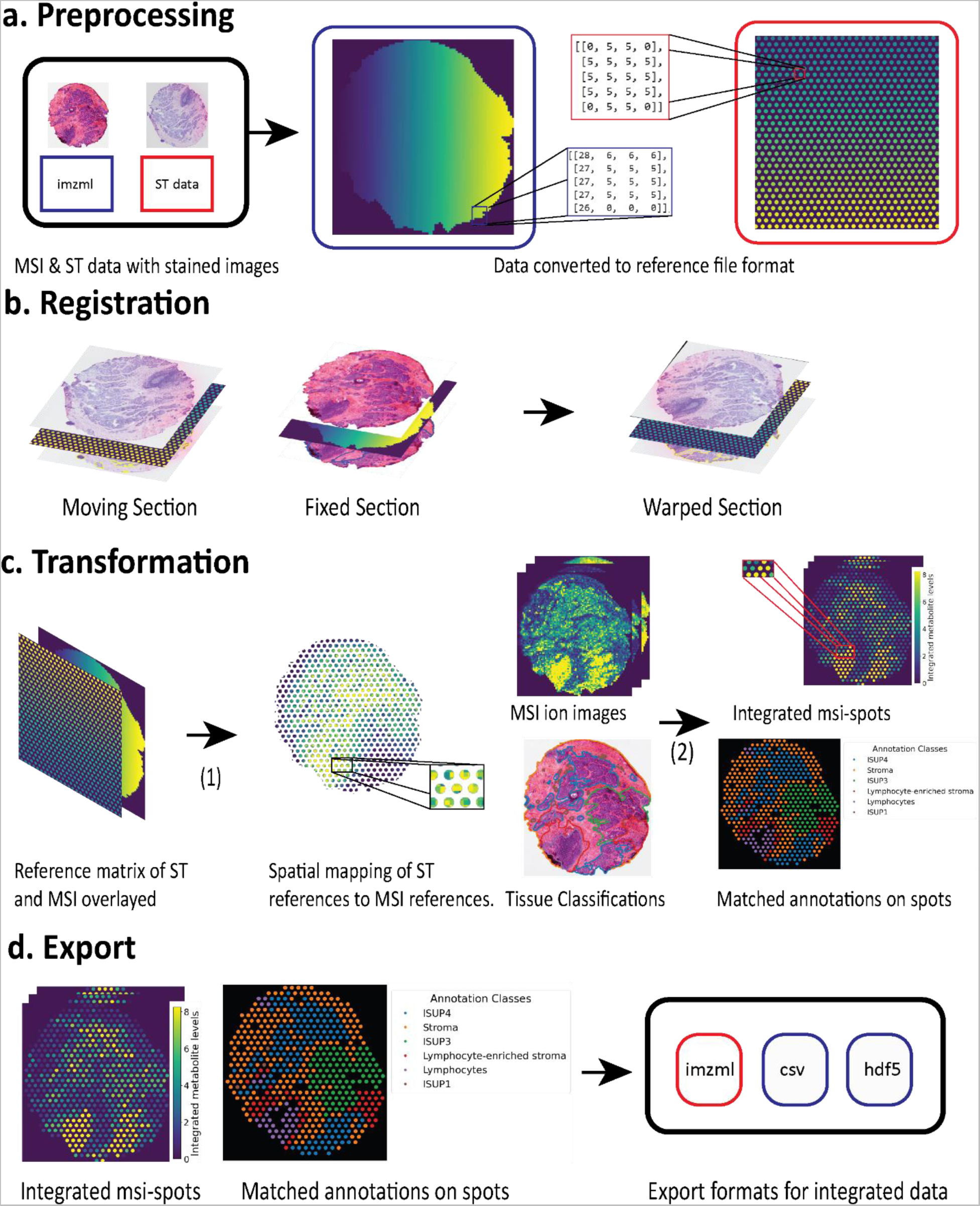
Integration workflow of MIIT. **a) Preprocessing.** Preprocessing of ST and MSI data. File formats are processed to reference matrices and registered to stained images, if necessary. Each number in a reference matrix is either a reference to on-tissue molecular data or 0 which denotes background. In this example, ST contains 2077 different spot references, MSI contains 7527 different pixel references to molecular data points. Different values are highlighted in different colors. **b) Registration.** Then the ST-section is registered to the MSI-section based on stained images. **c) Transformation.** (1) Reference matrix of ST is used to group MSI-data within the same spot regions and (2) grouped MSI-data is aggregated within each spot resulting in MSI-spots. If additional annotations are provided, integrated spots can be matched against these annotations as well. **d) Export.** Lastly, integrated spots are exported into the relevant file formats.

### Features & Extendibility

Although the main goal is to offer an efficient pipeline for the integration of spatial omics data, MIIT contains additional functions for processing spatial omics data and supports additional data formats such as pointset data, geojson and tissue annotation masks that can be included in the integration process. Moreover, additional spatial data types and different types of spatial omics can be easily added by implementing interfaces that are provided in MIIT. Although we focus on the integration of ST and MSI, MIIT can also be used for implementing different integration workflows. We recommend using the modality with the highest spatial resolution as the source section and the modality with the lowest resolution as the target section.

### GreedyFHist registration outperforms alternative method

The registration of serial sections is the centerpiece of MIIT’s integration pipeline. Therefore, we evaluated GreedyFHist’s registration accuracy on our fresh frozen prostate tissue samples. The evaluation of our tissue segmentation algorithm demonstrated a higher segmentation accuracy than a based alternative based on Otsu’s thresholding (Supplementary Results).

First, we investigated the registration accuracy of GreedyFHist on 212 pairs of adjacent serial sections (distance of 10 – 20 µm) by calculating the target registration error (*TRE*) between manually placed pairs of histologically distinct landmarks (Figure 4c, d; average number of landmarks per section: 77). The 212 pairs consist of images with 4 different types of staining (Figure 1). We compared GreedyFHist with HistoReg as both algorithms use the *Greedy* algorithm to compute the registration between preprocessed images and because HistoReg was the best open-source registration algorithm in the grand challenge for automatic non-rigid image registration (ANHIR)^8^. For each registration we first calculated the median TRE of all landmark pairs. To evaluate the accuracy over the whole dataset, we then calculated the median of median-TRE (MM-TRE) and the average median-TRE (AM-TRE). GreedyFHist resulted in significantly lower median-TRE (MM-TRE: 20.988 µm vs. 25.369 µm, AM-TRE: 44.096 µm vs. 217.064 µm; p ≈ 1.722 x 10^-18^, one-sided Wilcoxon signed-rank test; Table 3, Figure 4a,c, Supplementary Table 9) than HistoReg. GreedyFHist required on average 137.35 ± 16.27 seconds per registration compared to 68.02 ± 5.66 seconds with HistoReg. These results suggest, that by applying our preprocessing method prior to image registration through Greedy, we can robustly register stained images of fresh frozen serial sections though at the cost of a longer runtime.

**Figure 4.**
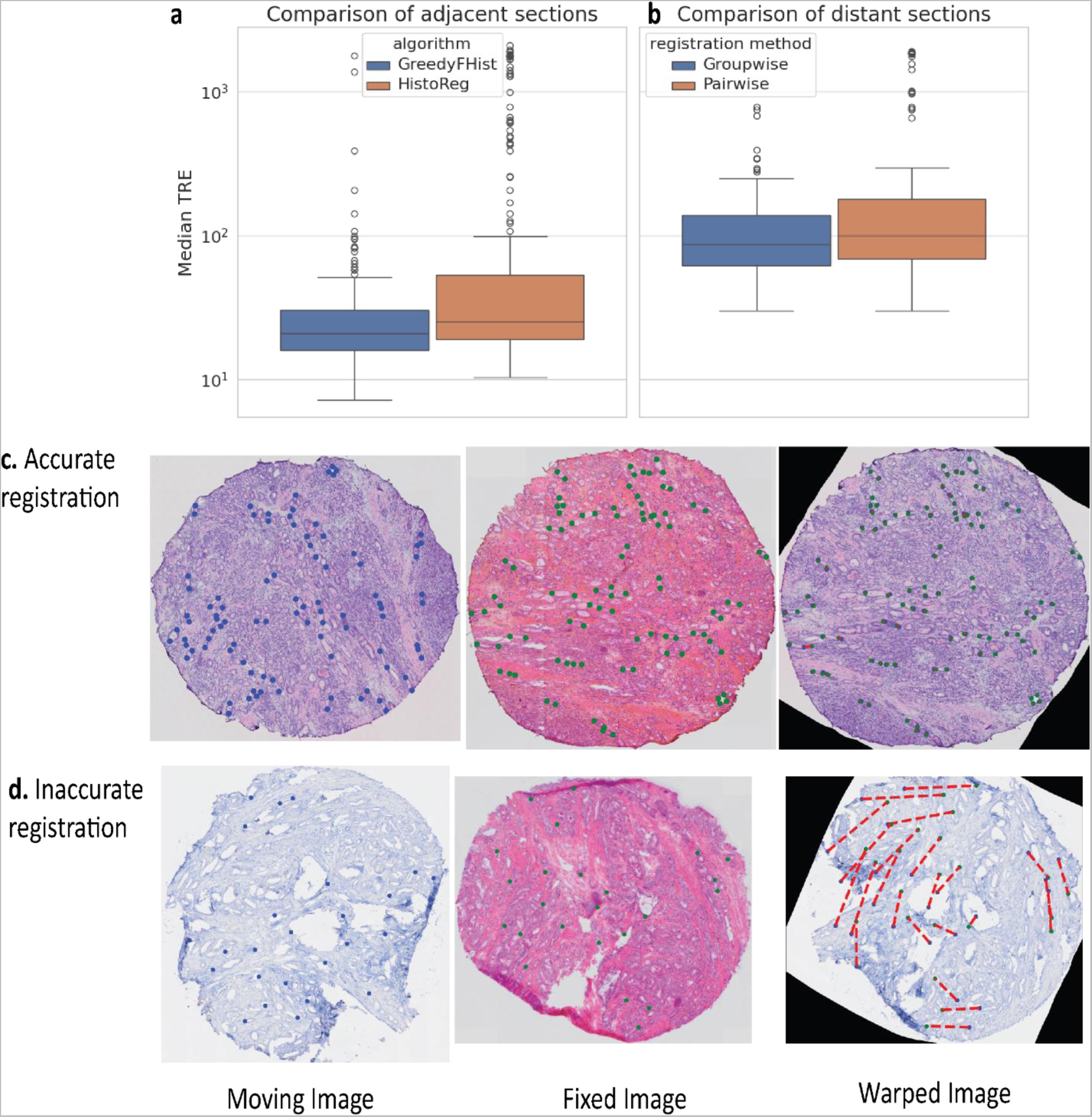
Assessing registration accuracy using landmarks. Comparison of error distribution for a) registration of adjacent sections between GreedyFHist and HistoReg and b) registration of distant sections between pairwise registration mode and groupwise registration mode using GreedyFHist. median-TRE is shown at a log-scale. Representative registration examples showing (c) an accurate registration (median TRE = 9.970 µm) and (d) an inaccurate registration due to tissue damage (median TRE = 593.006 µm). Landmarks of moving and warped landmarks are plotted in blue; landmarks of fixed images are green and the distance between warped and fixed landmarks for warped images is illustrated in a red dashed line.

**Table 3:** Registration performance of HistoReg and GreedyFHist on directly neighboring serial sections. Registration accuracy is reported as the median of the median TRE (MM-TRE), average of the median TRE (AM-TRE). AM-TRE metrics are reported with standard deviation (SD). TRE-based metrics are reported in µm, duration in seconds.

| Algorithm | MM-*TRE* | AM-*TRE* (± SD) | Duration (mean ± SD) |
| --- | --- | --- | --- |
| GreedyFHist | 20.988 | 44.096 ± 155.02 | 137.35 ± 16.27 |
| HistoReg | 25.369 | 217.064 ± 486.84 | 68.02 ± 5.66 |

### Groupwise registration improves registration of distant neighboring section

In multi-omics experiments, it may be necessary to register stained images that are not adjacent neighbors, but distant to each other. For instance, for the experimental setup depicted in Figure 1, ST was applied on section 2 and MSI on section 6 and 7 (which is equivalent to a distance of 40 – 100 µm). This motivated us to investigate how well distant sections can be registered with GreedyFHist. Therefore, we chose to register the 122 image pairs from sections with a distance of 5 sections which corresponded to 50 – 100 µm. Images that GreedyFHist could only register poorly during the pairwise evaluation (median-TRE > 200 µm; Figure 4d) were not included. We decided to analyze how well two distant images can be registered by evaluating two different strategies: *Pairwise registration*: Moving and fixed image were registered directly. *Groupwise registration*: Intermediate images were used to build a registration sequence between the moving and the fixed section.

The groupwise registration resulted in significantly lower TRE-metrics than the pairwise registration (MM-TRE: 89.871 vs. 100.406; AM-TRE: 176.813 vs. 292.452; p ≈ 0.015 for median-TRE, Table 4, Figure 4b). The higher TRE-metrics for the direct registration are expected due to the registration having to account for stronger effects of tissue heterogeneity whereas the groupwise registration adjusts to the changing tissue heterogeneity stepwise with each intermediate registration. Moreover, the higher TRE-metrics between distant and adjacent tissue sections are also expected due to stronger effects of tissue heterogeneity in distant sections compared to adjacent sections. To conclude, this experiment shows that to overcome the issue of registration of distant images, a groupwise registration strategy yields significantly better results than aiming to register images directly.

**Table 4:** Comparison of different strategies for registration of distant serial sections. Registration accuracy is reported as the median of median TRE (MM-TRE), average median TRE (AM-TRE). AM-TRE metrics are reported with standard deviation (SD). TRE-based metrics are reported in µm, duration in seconds.

| Registration method | MM-TRE | AM-TRE (± SD) | Duration (mean ± SD) |
| --- | --- | --- | --- |
| Distant pairwise registration | 100.406 | 292.452 ± 491.220 | 143.71 ± 15.482 |
| Distant groupwise registration | 89.871 | 176.813 ± 300.132 | 322.12 ± 43.721 |

### Spatial multi-omics and its integration using MIIT is capable of reproducing known biological associations

To demonstrate the applicability of MIIT to gain multi-omics biological insights, we choose to investigate a well-known metabolic mechanism of the prostate; citrate and spermine secretion. Prostate luminal cells secrete large amounts of the metabolite citrate into the prostate lumen. The high production of citrate is proposed to be the result of high zinc levels, which inhibits the citric acid cycle causing accumulation of citrate. Spermine is another metabolite which is secreted at high levels by the prostate and is highly correlated with citrate levels^23,28,29^. Secretion of citrate and spermine is lost during progression to high-grade prostate cancer and is absent in the prostate stroma^30,31^. Thus, high levels of citrate, zinc and spermine are a hallmark for normal prostate glandular tissue^32,33^. Our research group has previously developed a gene signature for citrate secretion gene signature (CSGS) in the prostate^23^, which links gene expression to metabolic citrate secretion. To demonstrate the potential of MIIT, we investigated the correlations between single sample Gene Set Enrichment Analysis (ssGSEA) scores^24^ calculated from CSGS on ST data and citrate, zinc and spermine levels from MSI^34^.

For investigating the relation between CSGS and metabolite levels, we chose a subset of 7 cancer-free tissue samples from different prostate cancer patients. For computing ssGSEA scores for CSGS we found 109 of 150 genes in CSGS in our ST data (Supplementary Table 8). ST and MSI in positive and negative ion mode were integrated using MIIT as described in the method section with an average MM-TRE of 86.04 µm (range: 38.17 µm – 139.22 µm; Supplementary Table 3). As illustrated in Figure 1, ST was performed on section 2 (ST-section), MSI in positive ion mode on section 6 (MSI-POS-section) and MSI in negative ion mode on section 7 (MSI-NEG-section). Spatial integration of these data resulted in two sets of MSI spots per sample, one for positive ion mode (MSI-POS-spots) and one for negative ion mode (MSI-NEG-spots). Citrate and zinc measurements were taken from integrated MSI-NEG-spots and spermine measurements from integrated MSI-POS-spots. Furthermore, histopathology annotations from ST-sections, MSI-POS-sections and MSI-NEG-sections were used to classify each ST-spot, MSI-POS-spot, and MSI-NEG-spot as either gland or stroma. To account for batch effects that can occur in different samples, we analyzed each sample separately. Integrated spots that did not cover tissue regions or were not classified as gland or stroma were excluded from further data analysis.

Differential expression analysis of samples comparing gland and stroma spots predominantly showed significantly higher CSGS score and higher citrate, zinc and spermine levels in glands (Mann-Whitney U rank test, Figure 5a-d; Supplementary Figure 13, Supplementary Table 4). Further, the CSGS score showed significant correlations with citrate, zinc and spermine for most samples (Figure 5e-h for one sample, Supplementary Figures 5,6). This confirms that the well-known prostate mechanisms are present in this multi-omics dataset and that CSGS, citrate, zinc and spermine are good candidates for a proof-of-concept demonstration of MIIT^30,31^.

**Figure 5.**
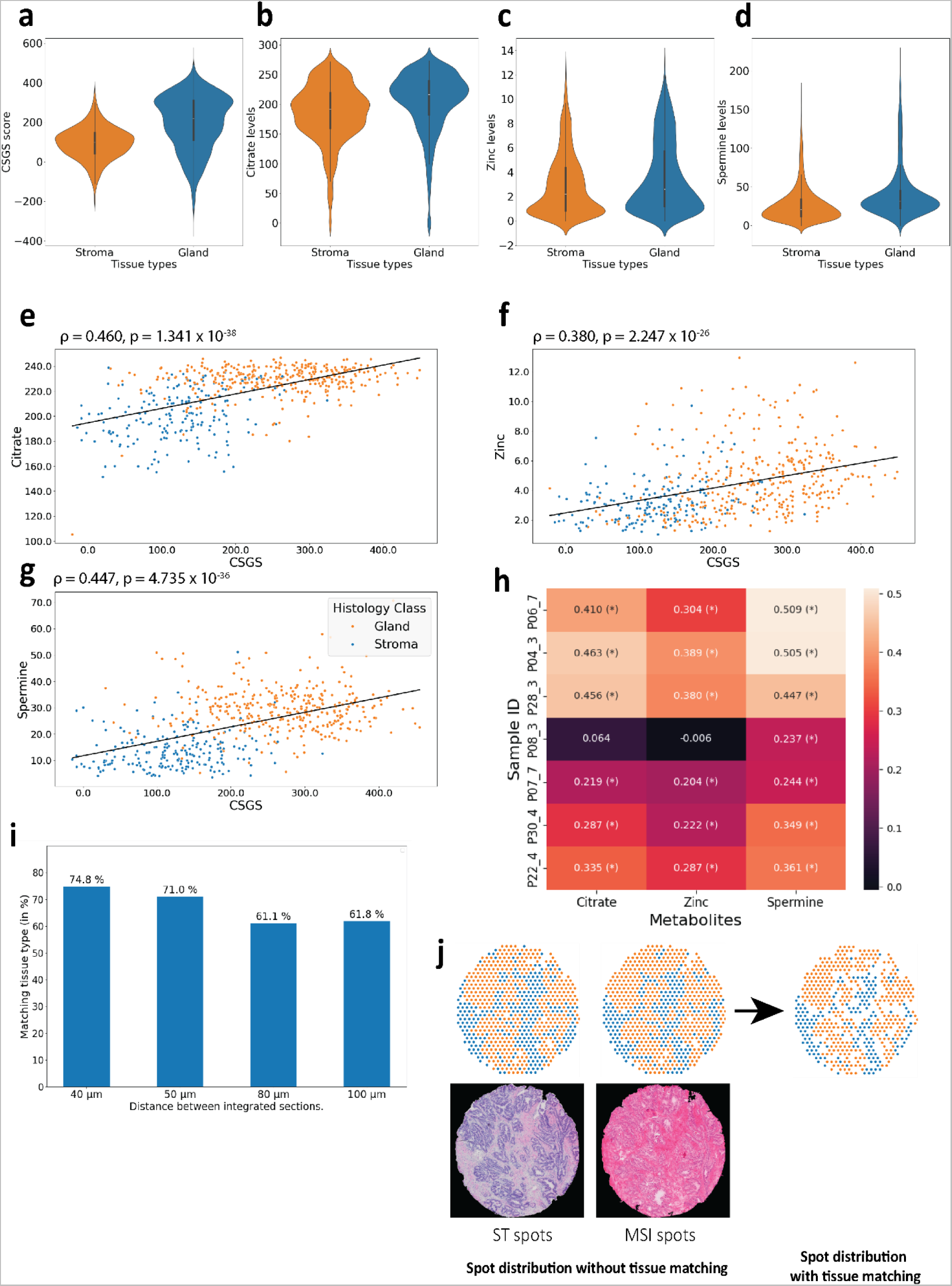
Comparing gene signature and metabolites between gland and stroma spots. Gene scores and metabolite levels in stroma and gland spots for a) GSCS, b) citrate, c) zinc, and d) spermine. e) Citrate, f), zinc, and g) spermine levels plotted against CSGS score for one sample (P28_03) for integrated spots colored according to tissue type. Linear regression lines, spearman correlation coefficient ρ and p-value are shown. h) Sample-wise correlation coefficients between CSGS and citrate, zinc, and spermine. (*) denotes significance (< 0.05). i) Percentage of successfully integrated spots after tissue type matching across different section distances. j) Distribution of gland and stroma spots in ST section and MSI section without tissue type matching and after tissue type matching. Spots that could not be assigned to either stroma or gland or had a different histopathology classification were discarded beforehand.

#### Tissue type matching improves correlation coefficients

Despite a high integration accuracy, image registration is not able to account for all morphological changes in tissue composition between neighboring sections. For the 7 samples used in this analysis, distance between the ST-sections and the MSI-sections ranged from 40 to 100 µm due to their sectioning order (Figure 1) and varying number of discarded sections. This results in some areas where ST-spots and MSI-spots have a different tissue type (Figure 5j). Therefore, we were interested in analyzing the effect of tissue heterogeneity between ST-section and both MSI-sections and whether correlation analysis are impacted by this. From the initial integrated datasets (termed **IntWithoutMatchHist**) we removed all spots with non-matching tissue types (termed **IntMatchHist**) which reduced the dataset by 34.7% for matching tissue types between ST-section and MSI-NEG-section and 32.72% between ST-section and MSI-POS-section (Figure 5i). Spearman correlations analysis between the CSGS score and metabolite levels demonstrated that only including histology matched spots increased correlations significantly for both citrate (*ρ̄* = 0.433 vs *ρ* = 0.297, p = 0.011, paired t-test on z-transformed ρ) and zinc (*ρ̄* = 0.330 vs *ρ* = 0.247, p = 0.046) (Figure 6a-d (1,2), Supplementary Table 4, Supplementary Figures 5, 6). For spermine, using matched histology spots had less impact with only nearly significantly higher correlation (*ρ̄* = 0.441 vs *ρ* = 0.361, p = 0.056). The reason for this could be that the distance between ST section and MSI-POS section is shorter than the distance between ST section and MSI-NEG section resulting in less difference in expression between CSGS and spermine. Though not above the significance threshold overall, we note that for some samples ρ of CSGS and spermine improved considerably (P06_7, P03_4; Figure 6d, Supplementary Table 5). These results show that taking changing histology into account when integrating serial sections is important when investigating tissue type specific biological mechanisms.

**Figure 6.**
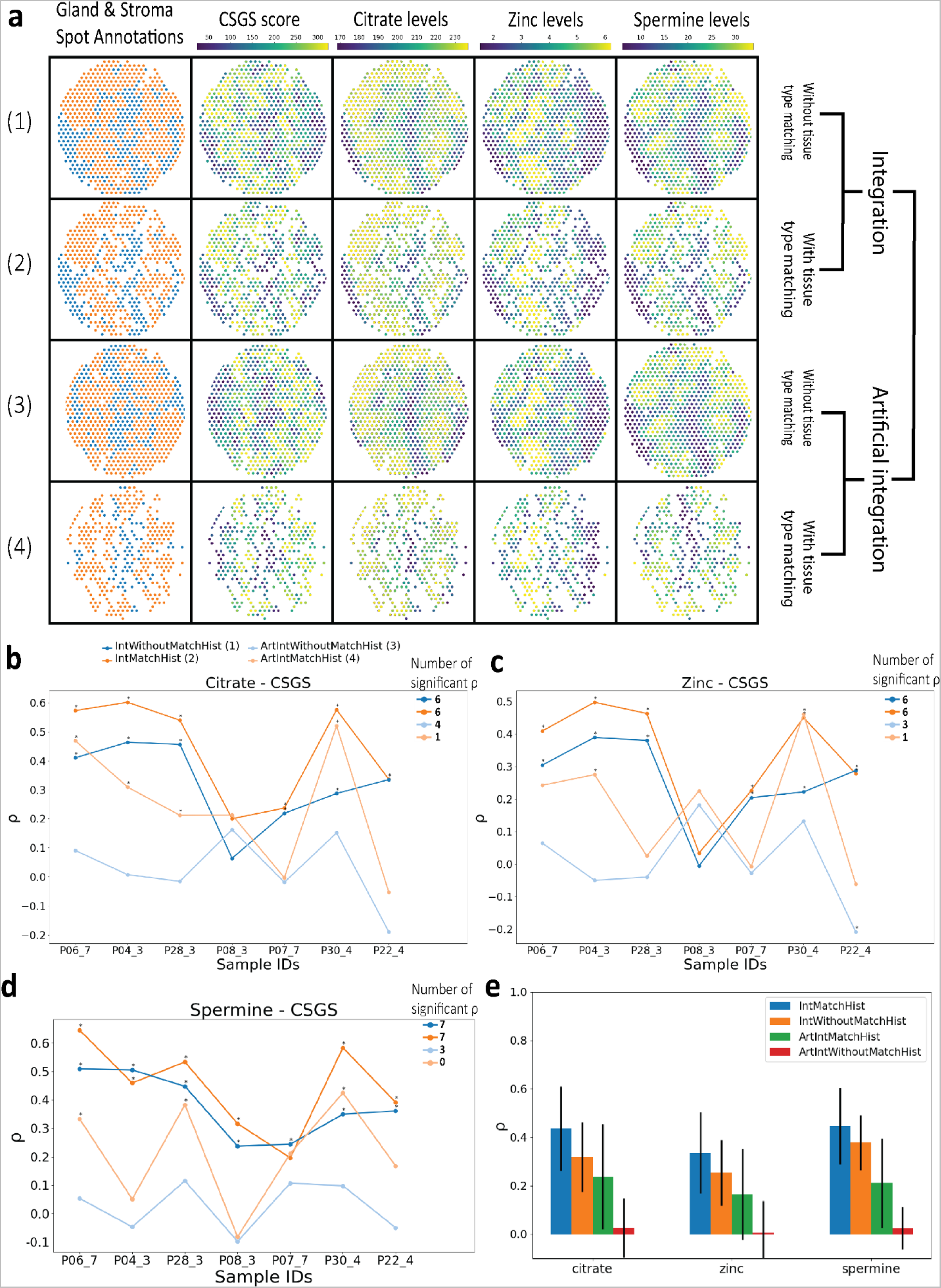
Comparison of different integrated spatial multi-omics datasets. a) Spot-wise distribution of gland and stroma, CSGS, citrate, zinc, and spermine for four different datasets for sample P28_3. b-d) Spearman correlation coefficient distribution for each sample for all 4 datasets. * denotes significant correlations (p < 0.001). e) Average ρ for each metabolite compared to the CSGS score. Error bars represent standard deviation.

#### Correctly integrated data performs better than artificially integrated data, demonstrating proper integration is important

To test the robustness of MIIT, we compared the IntWithoutMatchHist and IntMatchHist with dataset featuring deliberately poor integration. This artificially integrated dataset was created by rotating the ST-sections after registration by 180 degrees followed by integration, which is termed **ArtIntWithoutMatchHist**. Additionally, we performed tissue type matching on the artificially integrated data, creating **ArtIntMatchHist**, to investigate whether poorly integrated spots with the same tissue type give us similar results as correctly integrated spots. Not surprisingly, a much larger proportion of non-matching spots had to be degraded when the data was artificially integrated (Figure 6a, Supplementary Figure 2, 7-12). We evaluated all four datasets by comparing the mean spearman correlation coefficient ρ and the number of significant correlations. For both criteria for all three metabolites IntMatchHist performs best, followed by IntWithoutMatchHist, ArtIntMatchHist and ArtIntWithoutMatchHist (Figure 6b-e, Supplementary Table 6, Supplementary Figure 3). Interestingly, both correctly integrated datasets showed better results than the artificial ones showing that an accurate registration is necessary. The fact that IntWithoutMatchHist performed better then ArtIntMatchHist demonstrates that there is biological heterogeneity between spots of the same tissue type (gland and stroma areas). In other words, poorly integrating e.g. one gland spot with a different distant gland spot can weaken the biological interpretation of the data.

## Discussion

In this paper, we introduce MIIT (Multi-omics Imaging Integration Toolset), a novel and flexible framework designed to integrate various spatial-omics data. Using MIIT we defined a semi-automated workflow that merges spatial transcriptomics (ST) and mass spectrometry imaging (MSI) data into a combined dataset. It is customizable and can be extended to other types of spatial omics data. Our proof-of-concept analysis demonstrated MIIT’s capability by integrating ST and MSI data to validate a bulk-generated gene signature for citrate and spermine prediction in prostate cancer^23,29–33^. A key component of MIIT’s integration workflow is the registration of neighboring tissue sections using a novel non-rigid registration algorithm, GreedyFHist. We evaluated GreedyFHist on fresh frozen tissue samples with four different types of staining, achieving a high accuracy. Both MIIT and GreedyFHist are available as open-source software to use for all types of tissues.

GreedyFHist utilizes *Greedy* for computing affine and nonrigid registration parameters, similar to another state-of-the-art registration algorithm, HistoReg. However, by implementing a novel preprocessing pipeline GreedyFHist outperforms HistoReg’s registration accuracy (MM-TRE: 20.988 µm vs. 25.369 µm; AM-TRE: 44.096 µm vs. 217.064 µm; Table 3, Figure 4a). GreedyFHist includes background segmentation to remove noise and center the registration on the tissue region of interest, followed by mean shift filtering to denoise features in tissue images while preserving tissue boundaries. We further compute the center-of-mass to improve initial alignment for the affine registration. These additional steps contribute to GreedyFHist’s higher accuracy compared to HistoReg. Based on the registration of two stained images, we developed a groupwise registration method that leverages intermediate stained images to accurately register stained images up to 100 µm apart. This makes it possible to accurately integrate spatial omics data distributed over several serial sections. GreedyFHist interfaces with the well-known software QuPath, a digital pathology platform and is available as a separate software package. Therefore, it is also possible to use GreedyFHist outside of MIIT, e.g., for 3D image reconstruction. For the transformation from the spatial organization of MSI to the spatial organization of ST data, we calculated weighted statistics between each ST spot and the spatially matching MSI data. This approach is appropriate given the high-resolution dense representation of MSI data and the low-resolution sparse distribution of ST data. However, for use-cases in which data is mapped from low- to high-resolution (e.g., ST into MSI), advanced methods such as deconvolution techniques could be included in the integration workflow. Future research could explore deconvolution methods that directly work on integrated molecular data.

Despite growing interest in spatial multi-omics analysis, few computational open-source end-to-end pipelines exist for integrating various types of spatial-omics data. For instance, Sun et al.^3^ performed spatial integration between ST and MSI data by manually registering MSI data onto a neighboring HE-stained image of the ST section. They used MassImager Pro^TM^ for metabolomics and SCiLS Pro 2018b (Bruker Daltonics) for lipidomics. Unlike MIIT’s automated workflow, there is manual making it challenging to apply to large-scale datasets and neglects the changing tissue morphology between serial sections. Ravi et al.^2^ integrate MSI with ST data by registration of neighboring stained serial sections to map MSI to ST data followed by averaging MSI pixels that share the same ST spot coordinates. However, they rely on an affine registration method^35^, known to produce higher target registration errors than non-rigid registrations^7–9^ and have not evaluated their method on distant sections. In contrast, MIIT employs GreedyFHist to achieve high accuracy through non-rigid registration and has successfully integrated ST and MSI at a distance of 100 µm. An end-to-end pipeline similar to MIIT is the SMOx pipeline presented by Zhang et al.^4^, which they used to integrate ST data and lipidomics via MSI in prostate cancer samples. They performed non-rigid registration of serial stained images to align MSI and ST data. For the transformation of MSI onto ST data, SMOx used granularity matching with a Gaussian approach, whereas MIIT used weighted statistics over the shared area between each ST spot and MSI pixels. However, the biggest difference between the 2 pipelines is that SMOx is a proprietary software, while MIIT’s framework is open-source and customizable, providing users with more flexibility. An alternative sample preparation protocol by Vicari et al.^36^ enables the retrieval of MALDI MSI and ST from the same tissue section, circumventing the issue of registration of serial sections. Although this strategy limits the number of spatial omics types that can be used, it can be combined with MIIT by adding other types of spatial omics data to serial sections. Through MIIT’s customizable open-source design and the high registration accuracy provided by GreedyFHist, it is a viable alternative framework for spatial multi-omics integration.

Another excellent advantage of MIIT is its ability to extend seamlessly to other types of spatial omics with minimal prerequisites. This is achieved through grid projection into a standard reference matrix format, enabling spatial matching and non-rigid transformation of all spatial omics data with high accuracy. The registration between spatial omics data is performed by registering stained images, making the process independent of the omics methodology attached to each stained image. Molecular measurements are accessed only during the transformation step for mapping between different spatial-omics types. The only prerequisite is that each spatial omics data requires a reference image for registration. Although we used stained histology images as references, other image types could also be used, presenting a topic for future research. The easiness of adding additional types of spatial omics makes MIIT a useful framework for exploring new types of spatial-omics integration.

Nonetheless, GreedyFHist and MIIT have some limitations that we are addressing. Integrating multiple serial sections presents the challenge of unavoidable spatial alteration of histology. We found that including only integrated spots with consistent histology types between tissue sections significantly improved correlations between known associations of the citrate-spermine gene signature and the metabolites citrate, zinc, and spermine in prostate cancer tissue. However, this came at the cost of losing integrated spots. This loss was acceptable in our dataset due to the abundance of stroma and gland spots, but it could be problematic for smaller tissue components (e.g., perineural invasions on prostate tissue). Ensuring that molecular imaging experiments are performed on tissue sections as close as possible is crucial to reduce the effects of tissue heterogeneity, a problem not unique to MIIT. Another limitation is the occurrence of tissue damage^8,14^. Although GreedyFHist achieved high accuracy in registration, some images could not be registered accurately due to tissue damage (Table 3, 4, Figure 4), which can become problematic during groupwise registration. Developing methods to estimate registration quality and filter out unregistrable images could address this issue. These limitations affected MIIT directly since it required accurate registration for spatial multi-omics integration. MIIT addresses this issue through its flexible design, which allows us to exchange GreedyFHist’s registration with an alternative registration algorithm during registration. This alternative registration algorithm registers images using manually chosen landmarks (e.g. through external tools like Fiji^37^). MIIT has extra functionality to add custom registration algorithms easily to the integration pipeline making it easily adaptable to specific use-cases, e.g., for faster registration or different image modalities.

MIIT’s workflow is versatile, making no assumptions about the origin of the underlying data, making it valuable for a wide range of heterogeneous disease types. MIIT advances the investigation of the tumor microenvironment through multi-omics integration of spatially resolved transcriptomics and metabolomics^38–40^, enabling comprehensive computational analysis of different classes of molecules and is useful for all types of tissues, not only cancer tissue. Future work will focus on analyzing the spatial relationship between molecules detected by different omics methods within the prostate tumor microenvironment to reveal insights into prostate cancer aggressiveness. Additionally, MIIT’s open and flexible design allows for future developments, such as adding automated tissue annotations^41^ and integrating other types of molecular imaging data. MIIT could also be used to integrate spatial-omics methods applied to whole organ tissue sections with MRI from the same organ, requiring algorithms capable of registering stained histology images with MRI data^42–44^. Establishing normalization methods for analyzing spatial multi-omics datasets is also necessary but was outside this work’s scope.

To conclude, we presented a novel framework for the integration of spatially resolved molecular imaging. MIIT is openly accessible and customizable to handle a variety of different types of spatial-omics data. We also developed an algorithm for the registration of tissue samples that achieved high accuracy on a set of fresh frozen tissue samples and is available for all tissue types. MIIT is implemented and released as a software package at *github.com/mwess/miit* along with documentation and tutorials whereas GreedyFHist can be found at *github.com/mwess/greedyfhist*.

## Methods

### ProstOmics dataset

All data used in this study is part of the ProstOmics dataset, from our larger project “’Tissue is the issue’: a multi-omics approach to improve prostate cancer diagnosis” (ERC: 758306). Parts of this project have been used in previous publications of our group^18,45^, but the set-up for this study is presented in Figure 1. Several parts of this dataset have been used in previous work of this group: HE and MTS stained sections, histopathology on HE stained sections and ST data have been used by Andersen et al.^18^. ST data has been used by Kiviaho et al.^20^. ST (including HE staining) and MALDI-TOF MSI data in negative ion mode (including HES staining) has been used by Krossa et al.^19^.

### Patient inclusion and sample collection

The regional ethics committee of Central Norway approved this research (identifier 2017/576). All methods were carried out according to national and EU ethical regulations. Human prostate tissue samples were collected after informed written consent was given by prostate cancer patients undergoing radical prostatectomy. Samples of eight patients, who were not treated prior to surgery, were collected from St. Olav’s hospital, Trondheim, Norway between 2008 and 2016. Three patients were considered relapse-free as no confirmed relapse occurred after 12 years and five patients were classified as relapse as metastasis occurred within three years after surgery. A 2 mm thick slice was cut from the middle of the prostate (transverse plane), snap frozen and stored at −80°C as described by Bertilsson et al.^46^immediately after surgery by expert personnel at Biobank1®, St Olavs University Hospital, Trondheim, Norway. A range of 8 – 13 tissue samples (3 mm in diameter) were collected from each fresh frozen tissue slice, using an in-house built drill system. Based on HES-stained tissue annotations 4 sample cores were selected from each patient’s tissue slice with 2 samples containing cancer tissue, one sample being cancer tissue adjacent and one sample from a far away region on the tissue slice containing no cancer tissue, giving a total of 32 samples.

### Cryosectioning

Tissue sections from each tissue sample were cut with 10 µm thickness inside a cryostat at −20°C (Cryostar NX79, Thermo Fisher Scientific). For this study, we selected sections (1-3, 6-11) from various staining methods presented in Figure 1. In total, we used 288 sections from all 8 patients where 9 sections were collected from each tissue core. Four conductive slides were vacuum packed and stored at −80°C until further use.

### Staining methods

All sections were stained for various experiments with Hematoxylin and Eosin (HE; n = 32), Hematoxylin, Erythrosine and Saffron (HES; n = 160), Immunohistochemistry (IHC; LPS and LTA; n = 61; Supplementary Methods) and Masson Trichrome Staining (MTS; n = 32) and scanned at 20x magnification. We note that section 3, 6 and 7 were stained after performing MALDI-TOF MSI (n = 68). Additionally, HES-stained tissue on section 3 (n = 32) was heated for 5 minutes at 95°C before MALDI-TOF MSI according to the protocol developed by Høiem et al.^45^. For each core, sections were stained in the same order (see Figure 1).

### Histopathology

HE scans from section 2 and HES scans from section 6 and 7 were independently evaluated and annotated by two experienced uropathologists (T.V. and Ø.S.) using QuPath version (v 0.2.3) and (v 0.4.4). The identified areas were lymphocytes, stroma, epithelial areas, glands, and tissue borders in addition to cancer areas according to the Gleason Grade Group system. Furthermore, a consensus pathology evaluation was reached in agreement with both pathologists. For each ST spot from section 2, the fraction of different tissue types and regions present were calculated.

### Spatial transcriptomics

Sequencing libraries were created from the tissue sections by using the Visium Spatial Gene Expression Slide & Reagent kit (10X Genomics) following the manufacturers manual. In brief, tissue sections were fixed using methanol, followed by HE staining, and scanning of slides at 20x magnification. For microscopic scanning a coverslip was put over the sections and removed afterwards. To capture mRNA, tissue sections were incubated with permeabilization enzyme for 12 minutes, which previously had been optimized using the Visium Spatial Tissue Optimization Slide & Reagent kit (10X Genomics). A second strand mix was added to create a second strand, followed by amplification of cDNA by real time qPCR. The amplified cDNA library was quantified with qPCR using the QuantStudio™ 5 Real-Time PCR System (Thermo Fisher) and the cDNA libraries were stored at −20°C until further use. Paired-end sequencing was performed on an Illumina NextSeq 500 instrument (Illumina®, San Diego, USA) using the NextSeq 500/550 High Output kit v2.5 (150 cycles).

### MALDI-TOF MSI

For MALDI-TOF MSI, 2 sections from each core were used, with position 6 for positive ion mode and position 7 for negative ion mode, resulting in a total of 64 sections. Before matrix application, all vacuum-packed slides with tissue sections were left on the benchtop for at least 20 minutes before opening the vacuum-bag. Two different matrixes, 2,5-dihydroxybenzoic acid (DHB) and N-(1-naphthyl) ethylenediamine dihydrochloride (NEDC), were prepared by dissolving DHB in 70% methanol/0.1% trifluoroacetic acid (concentration 20 mg/mL) and NEDC in 70% methanol (7 mg/mL). The HTX TM-Sprayer™ system was used to spray the matrix onto the tissue sections, with 14 and 18 layers of matrix for DHB and NEDC, respectively. The rapifleX™ MALDI Tissuetyper^TM^ (Bruker Daltonics) equipped with a 10 kHz laser was used to measure all tissue sections, shooting 200 shots per pixel at a 10 kHz frequency with a spatial resolution of 30 μm. Prior to all measurements, the instrument was calibrated using red phosphorus. Tissue sections covered with DHB matrix were measured in positive ion mode with a mass range of *m/z* 100–1000, while tissue sections covered with NEDC matrix were measured in negative ion mode with a mass range of *m/z* 40–1000. All measurements included separate matrix only regions. After data acquisition, the slides were stored at 4°C until staining with hematoxylin, erythrosine and saffron (HES). The MSI data were first imported into FlexImaging (Version 5.0, Bruker Daltonics) where we applied binning (factor 0.8). Then, MSI data were imported into SCILS Lab (Version 2024a Pro), baseline corrected using top-hat, normalized using root-mean square and quantified intensities as max peak height. We selected masses that we identified in a previous publication^47^. Tissue borders were annotated using the “Create new polygonal region” function. MSI data were exported in imzML^48^ along with tissue border annotations in SCILS’s srd format. For this study, we extracted the mass intensity of citrate (m/z 191.0 [M-H]^-^) and zinc in the form of ZnCl^-^ (m/z 174.8 [M]^-^) from the negative ion mode data and spermine (m/z 203.2 [M+H]^+^) from the positive ion mode data.

### Preprocessing of stained histology images

Tissue masks that separate tissue and background were used as training data for training our YOLO8-based segmentation model. They were created in QuPath (v 0.4.4). A threshold was used on the average intensity signal of each pixel after a gaussian filter (sigma: 1) was applied. The thresholds for HE, HES and MTS were set between 208 and 215, for IHC the threshold was set between 218 and 225. The threshold was manually adjusted on each tissue image if necessary. Tissue masks were manually corrected to remove wrongly labeled areas. Stained images, tissue masks and histopathology were exported using a groovy script (https://github.com/sekro/spatial_transcriptomics_toolbox) to *tiff images*. The resolution for each image was set to 1 pixel per µm.

### Hardware setup

All experiments were performed using the same hardware: Intel Core Processor Broadwell (32-core; 32 threads; Broadwell).

### Tissue Registration

GreedyFHist

Pairwise Registration:

#### 1. Segmentation

To remove background noise from the image and center the image around the tissue area, we performed background segmentation: Images were resampled to a resolution of 640×640 and converted to grayscale. Spurious noise from the background of the image was removed using a method by Chambolle^49^, that performs total variation denoising^50^. Then we used the YOLO8 model to perform tissue segmentation on the image. The model was trained for tissue segmentation on our dataset. From the segmented image, small classification artifacts (e.g. due to contamination on the slide) were filtered out by removing any region with a smaller area than 10000 µm^2^. The area threshold is dependent on the input data and can be adjusted by the user. The tissue mask was then resampled to back to its original size. Since the process of identifying correct tissue masks can vary between different experiments, we have implemented the option for users to supply their own tissue mask (e.g., from manual annotations) in GreedyFHist. This flexibility allows the user to apply masks for selected ROI on the tissue. After applying tissue masks to crop images, both images are padded to the same uniform shape.

#### 2. Denoising

Stained images are usually high-resolution images containing a large amount of detail. During registration such details may act as noise which negatively influences registration accuracy. Therefore, removing such noise while keeping major histological features intact is required. In the next step, we applied OpenCV’s implementation of mean shift filtering^51^ as a method of denoising: 1. Images are resampled to 512 x 512 pixels to improve filtering speed. 2. Images are converted to the HSV color space^52^. 3. Mean shift filtering requires a spatial and color window radius that determine the degree of filtering. We set these parameters for the spatial window radius to 20 and for the color window radius to 15. 4. Images are converted back to RGB color space and resampled to their original size.

#### 3. Grayscale conversion and downsampling

After denoising, images are smoothed with a Gaussian kernel (sigma: 2) to avoid aliasing effects during downsampling, resampled to a resolution of 1024 x 1024 pixel and converted to grayscale to further reduce image complexity. Images are then padded with 100 pixels on all sides to leave room for deformations during registration.

#### 4. Affine Registration

We performed an affine registration to find a global alignment between moving and fixed image. For computing the registration between two preprocessed images we use the *Greedy*, a registration tool that implements the greedy diffeomorphic registration algorithm (https://github.com/pyushkevich/greedy). *Greedy* has originally been developed for medical image registration, but has also used for the registration of stained images in a previous study^7^. We adapt some findings from this study, namely the use of the NCC kernel metric as a similarity measure and the postprocessing to rescale the computed transformation matrices to the image’s original resolution. We use *Greedy* in version 1.0.1, though other versions should be compatible with our registration pipeline as well.

Preprocessed images are exported in the nifty format using SimpleITK^53–55^. We initiated Greedy with a transformation that centers both images at their center-of-mass which we computed using the tissue masks. Then Greedy performs a random search to find a suitable rigid registration followed by a multi-resolution pyramidic approach for the affine registration: The registration is initially computed on downscaled images which is then recursively refined on upscaled images until the full resolution is reached. *Greedy* computes the registration using the limited-memory Broyden-Fletcher-Goldfarb-Shanno algorithm^56,57^ and we used the NCC kernel metric with a kernel size of 1% of the preprocessed image resolution (i.e. 10 pixels). Overall, we achieved the best result with the scaling factors 4, 2, and 1 though in some instances scaling factors 8, 4, 2, and 1 resulted in more accurate registrations.

#### 5. Non-rigid registration

After the affine registration, a nonrigid registration is performed to find a non-uniform transformation that can align the moving and fixed image locally which improves on the previous global registration. ^58^ The same parameters as for the affine registration were applied for the non-rigid registration. We additionally initialized the registration by supplying the computed affine registration and set two regularization parameters (pre-sigma: 6, post-sigma: 5). Table 5 contains all parameters passed to Greedy for the affine and non-rigid registration.

**Table 5.** Parameters that are used by *Greedy* to register preprocessed images.

|  | Affine registration | Non-rigid registration |
| --- | --- | --- |
| Resolution | 1024 x 1024 | 1024 x 1024 |
| Kernel metric + size | NCC, 10 | NCC, 10 |
| Pyramid iterations | 100, 50, 10 | 100, 50, 10 |
| Rigid search iterations | 10000 | - |
| Rigid search: standard deviation of rotation | 180 | - |
| Pre-sigma | - | 5 |
| Post-sigma | - | 5 |

#### 6. Postprocessing after affine and non-rigid registration

Tissue registration will result in two transformation matrices, one for affine and one for non-rigid registration. We rescaled both transformation matrices to match the original image’s sizes. The cropping and padding operations applied during preprocessing are expressed as translation transformations using SimpleITK^54,55^. These transformation matrices are composited along with the affine and non-rigid transformation matrices into a single displacement field matrix. This reduces interpolation artifacts during transformation as well as the number of image operations applied during transformation from moving to fixed image space.

#### 7. Transformation from moving to fixed image space

Using SimpleITK the displacement field is applied to warp image data from the moving to the fixed image space. GreedyFHist supports image data, image masks with annotations, pointset data in tabular form and pointset data in the geojson format. For image and pointset data, linear interpolation is used, and for annotation masks nearest neighbor interpolation is used.

### Registration Evaluation Measures

The registration performance of GreedyFHist was evaluated by using metrics based on the target registration error (TRE) for evaluating the accuracy between registered images. Through manual annotation of the stained sections, we set on average 77 landmarks of distinct histological features between any pair of serial section. There were 3 annotators in total and each set of annotations were cross validated. These metrics have previously been used in the ANHIR challenge for evaluating the accuracy of different registration algorithms^8,9^. The TRE is defined as

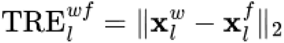

where 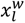 and 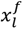 are a pair of matching landmarks *l* from warped and fixed landmarks. ||. ||_2_ denotes the euclidean distance. We evaluated each registration pair using the median TRE,

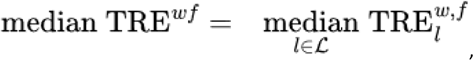

where 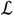 is the set of all landmarks between a pair of fixed and warped images. To evaluate the accuracy over the whole dataset we calculate the median of median-TRE,

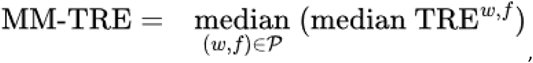

and the average median-TRE,

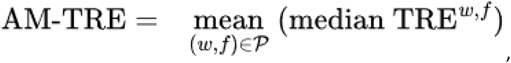

where 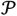 is the set of all registration pairs. Since all images are exported at a resolution of 1 µm per pixel, all errors are computed in µm.

### MIIT – Multi-omics Imaging Integration Toolset

#### 1. Preprocessing

The main purpose of this step is to set up easily generalizable data formats necessary for integration and to ensure a spatial alignment between stained and molecular imaging data.

##### Reference file format conversion

Spatial omics data is often stored in parametric data formats, e.g., ST-spots are circles with a center and a diameter, whereas MSI-pixels are described by a pixel location and resolution such that one pixel occupies one MSI spectrum. To allow for accurate non-rigid transformations, we performed a grid projection in which spatial omics data is translated into a reference matrix format. In this format, each spatial omics datapoint is represented by a reference and projected onto a matrix with the same resolution as the stained image.

##### Registration between stained image and molecular data

It is necessary to ensure that stained image and spatial omics data are spatially aligned before integrating can proceed. ST is preprocessed using 10x Genomics Space Ranger 1.0.0 which also guarantees an accurate registration between stained image and *ST-spots*. For the registration of MSI data to stained image, MIIT uses a registration method that is based on Verbeeck et al.^59,60^ (Supplementary Figure 4):

1. The HES-stained image is segmented using the YOLO8 based tissue segmentation method described in the preprocessing section of GreedyFHist and then cropped to the tissue’s boundary section.
2. A feature image for the MSI data is derived from the first principal component of the MSI data’s PCA spectrum. The feature image is rescaled to the stained image’s resolution (e.g., 1 µm per pixel).
3. Both images are padded symmetrically to create a uniform shape and then padded with 100 pixels to allow room for deformations. NiftyReg^61^ is used to find a rigid registration with the stained image used as the fixed and the MSI data as the moving image. We opted for a rigid registration, which only performs translation and rotation for image registration. This reduces deformation effects to the MSI-pixels, though options for affine and non-rigid registration can be selected as well. Alternatively, if segmentation masks for the stained image or MSI data are available, these can be used instead for the registration.

#### 2. Registration

The goal of this step is to register the source section with the target section. This is achieved by using GreedyFHist to register the stained images of the *source section* and the *target section*. The resulting transformation is then applied to all spatial data types in the source section.

In the case that GreedyFHist’s registration fails, e.g. due to tissue damage, we also provide an alternative registration algorithm based on scikit-image and manually provided landmarks (e.g. from Fiji^37^) (**Supplementary Methods**).

#### 3. Transformation

In this step, the spatially aligned data is used to transform between the spatial organization of the spatial omics data of the source and target section. For each data point in the target data space, the overlap with data points in the source data space is computed. We chose to use the modality with the lowest resolution (ST, 50 µm circular spots placed in a hexagon pattern 100 µm apart) as the target section and the modality with the finest resolution (MSI, 30 µm square pixels with no space between) as the source section. Each individual overlapping area in the source data space is then aggregated based on basic statistical descriptors. In practice, this means that the overlapping area between each *ST-spot* and the registered *MSI-pixels* is computed. Using the overlapping area, we can calculate the fraction that each *MSI-pixel* contributes to the shared area. Then we merge each metabolite intensity from the MSI space to the ST space by computing weighted statistical features (i.e., minimum, maximum, mean, standard deviation, median) over the shared area using the area fraction of each MSI-pixel as weight. We refer to the integrated MSI data as *MSI-spots*.

#### 4. Export

For the integration of ST spatial transcriptomics and MSI we export integrated data into a table format that uses barcodes from the ST count matrices as identifiers. Though other export options exist as well: Data can also be exported as an image stack with dimensionality n x w x h, where w and h refer to the dimensionality of the target sections image space and n refers to the size of each datapoint (e.g. the number of spectra in each MSI-spot). MSI integrated data can also be exported in the imzML format.

### Analysis of integrated spatial multi-omics dataset

After spatial integration, we applied tissue masks to all spots and filtered out any spots with less than 80% tissue coverage. Two samples (P08_3, P22_4) contained spots that were classified as “lymphocytes”. These spots were removed from the analysis. Correlations measured between CSGS and citrate, zinc, and spermine were computed using Spearman correlation.

## Author contributions

The manuscript was written by MW with help from MKA, MBT, SK and MR. MW designed and implemented algorithms and framework, SK provided scripts for exporting images and annotations from QuPath. MKA and SK consulted on data analysis, spatial transcriptomics and MALDI-TOF MSI. MR provided gene signature scores for CSGS. MW, EM and JC annotated serial sections with landmarks. ØS, TV and EM annotated tissue sections.

## Supporting information

Supplementary Results

## Acknowledgement

This research was funded by the European Research Council (ERC) under the European Union’s Horizon 2020 research and innovation program (grant agreement no. 758306), Norwegian University of Science and Technology (NTNU), the Liaison Committee between the Central Norway Regional Health Authority (RHA) and NTNU, the Norwegian Cancer Society and Terje Eugen Johnsen funds. All tissue samples were collected and stored by Biobank1, St. Olav’s Hospital. Tissue sectioning, staining and scanning were performed by or in collaboration with the Histology lab at the Cellular & Molecular Imaging Core Facility at NTNU. Transcriptomics experiments were carried out at the Genomics Core Facility at NTNU. MALDI MSI was acquired using instrumentation at the MR Core facility, NTNU.

## Data availability

The data can be provided by May-Britt Tessem pending scientific review and a completed material transfer agreement. Requests for access to the data should be submitted to.

