## Supplementary Methods for "Spatial Integration of Multi-Omics Data using the novel Multi-Omics Imaging Integration Toolset"

**Landmark-based registration**

This algorithm uses manually placed landmarks on the moving imagen and the fixed image. We find an affine registration based on landmarks using scikit-images^1^ estimate function. This function computes the affine transformation matrix using the least-squares method. Images and pointsets are transformed using linear interpolation whereas discrete annotations are transformed using nearest neighbor interpolation.

**Immunohistochemistry staining of LPS and LTA**

After fixation (4% formalin, 15 min), washing (2x TBS-T), and peroxidase inactivation with 0.3% hydrogen peroxide, cryo-sectioned tissue was blocked for 2h with 10% normal serum, 1% BSA in TBS at room temperature. Next, sections were incubated at 4°C over night with primary antibodies (anti-lipopolysaccharides (LPS), Abcam ab35654, 2,5 µg/ml in 1%BSA, TBS-T; anti-lipoteichoic acid (LTA), Thermo MA1-7402, 2 µg/ml in 1%BSA, TBS-T). After washing (2x TBS-T), secondary antibody, 3,3′-Diaminobenzidine (DAB), and hematoxylin staining were done according to manufacturer’s recommendations (EnVision anti-mouse-HRP/DAB+ system, Agilent). Digital images (bright field, 136,866 nm/pixel resolution) were obtained using an Olympus VS200 ASW 3.3 (Build 24382) slide scanner.

1.	van der Walt, S. *et al.* scikit-image: image processing in Python. *PeerJ* **2**, e453 (2014).
