## Supplementary Results for "Spatial Integration of Multi-Omics Data using the novel Multi-Omics Imaging Integration Toolset"

*Evaluation of tissue segmentation*

For the evaluation we used 285 stained images from the ProstOmics dataset for which we generated tissue masks as described in the methodology section. We assessed the quality of the YOLO8 based tissue segmentation with the metrics precision, recall, Dice similarity coefficient (DSC)^1,2^ and Hausdorff distance^3,4^. We validated our model using k-fold cross validation (k-fold; k=5) and an example can be seen in Supplementary Figure 1. In each fold, we separated our data into a training (80%) and test set (20%). We compared the results of our tissue segmentation algorithm to a baseline algorithm based on Otsu’s method^5^: We used the same processing techniques as with our tissue segmentation but replaced the prediction from the YOLO8 model with the results of Otsu’s method. Since Otsu’s algorithm does not require any training data, it is not necessary to perform cross validation.

Supplementary Table 1 summarizes the DCS, Hausdorff distance, precision and recall for the comparison between the YOLO8 based segmentation and the Otsu variant. The result shows that the YOLO8 based tissue segmentation outperforms the Otsu variant clearly by having a lower Hausdorff distance (15.736), a higher DSC (0.979) and recall (0.997). For the precision, our method performs slightly worse (0.982 against 0.991). For all metrics, our method achieved a lower standard deviation, signaling a higher stability of segmentation. The one-sided Wilcoxon signed-rank test showed significance for all metrics (p < 0.0001). Another point of interest is the runtime of our segmentation method as one of reasons for choosing the YOLO8 architecture was its capability of running inference efficiently on a CPU. To improve execution time of the YOLO8 based segmentation, we exported the trained model to the ONNX format which is designed to improve runtime of machine learning models^6^. We noticed a mean runtime of 1.06 seconds for segmenting one image using YOLO8 based segmentation and a mean runtime 0.02 seconds for one image segmentation using Otsu. Though Otsu’s method runs faster, we consider the runtime of our segmentation algorithm to be fast enough and negligible due to the long runtime typically associated with image registration.

**Supplementary Table 1: Averaged cross validation results for YOLO8 based tissue segmentation and Otsu segmentation**. Results are reported with mean and standard deviation.

|  | YOLO8 based tissue segmentation | Otsu segmentation |
| --- | --- | --- |
| Hausdorff distance (in µm) | 69.159 ± 54.434 | 140.814 ± 150.031 |
| DSC | 0.975 ± 0.021 | 0.924 ± 0.139 |
| Precision | 0.982 ± 0.020 | 0.990 ± 0.025 |
| Recall | 0.997 ± 0.007 | 0.933 ± 0.138 |

Next, we were interested in evaluating how robust our method performs on unknown staining. To emulate this scenario, we applied “leave-one-stain-out" (LOSO) validation where images of one staining type are removed from the training set and used as the test set. Consequently, each model is then evaluated on a type of staining that was not part of the training. We repeated this validation such that images from each staining method have been part of the test set. Table 2 summarizes the results of the LOSO validation. The highest performance for Hausdorff distance and DSC was achieved for Masson’s trichrome staining (MTS; 8.348, 0.989) in the holdout set and the worst performance for immunohistochemistry (IHC; 0.967, 24.465). We attribute this to the fact that MTS is a staining method that results in intense image features whereas IHC staining is less intense which is more challenging for the segmentation algorithm. Nonetheless, we consider robustness of our model for all staining methods as good enough for the purpose of tissue segmentation as a preprocessing step for tissue registration.

**Supplementary Table 2: Leave-one-stain-out cross validation separated for each staining method in the test set.** The staining methods are separated into hematoxylin & eosin (H&E), hematoxylin, erythrosine & saffron (HES), Masson’s trichome staining (MTS) and immunohistochemistry (IHC).

|  | Hausdorff distance (in µm) | DSC | Precision | Recall | Size of test set |
| --- | --- | --- | --- | --- | --- |
| H&E | 58.256 ± 57.009 | 0.983 ± 0.011 | 0.988 ± 0.010 | 0.997 ± 0.002 | 32 |
| HES | 65.731 ± 53.043 | 0.975 ± 0.023 | 0.980 ± 0.022 | 0.998 ± 0.004 | 160 |
| MTS | 36.382 ± 24.136 | 0.987 ± 0.008 | 0.991 ± 0.008 | 0.997 ± 0.001 | 32 |
| IHC | 105.082 ± 63.648 | 0.964 ± 0.022 | 0.971 ± 0.022 | 0.996 ± 0.007 | 61 |

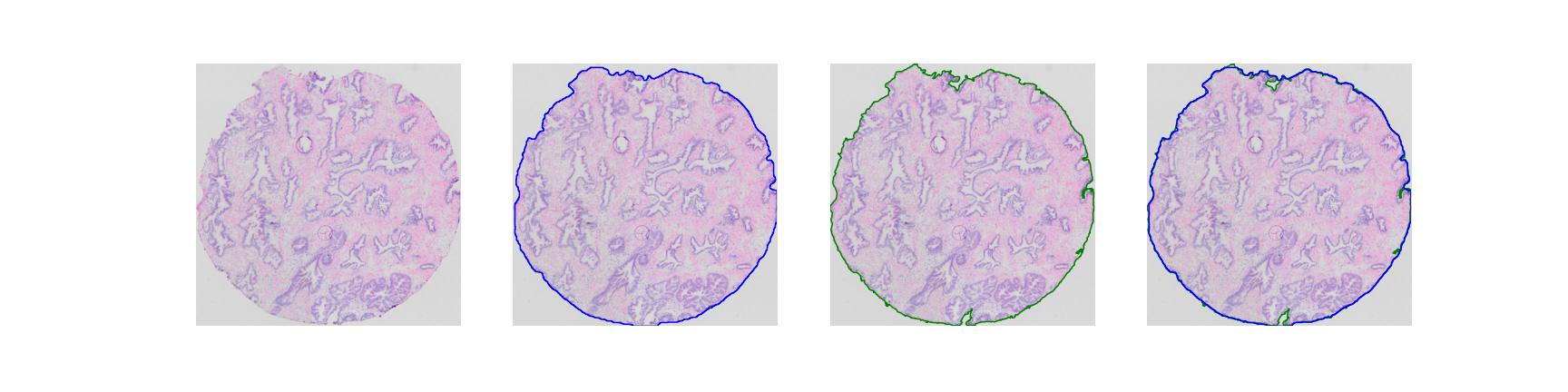

**Supplementary Figure 1. Validation of bounding box segmentation of stained image.** Blue highlights our segmentation method, green bounding box according to validation data.
