## Supplementary for "Spatial Integration of Multi-Omics Data using the novel Multi-Omics Imaging Integration Toolset"

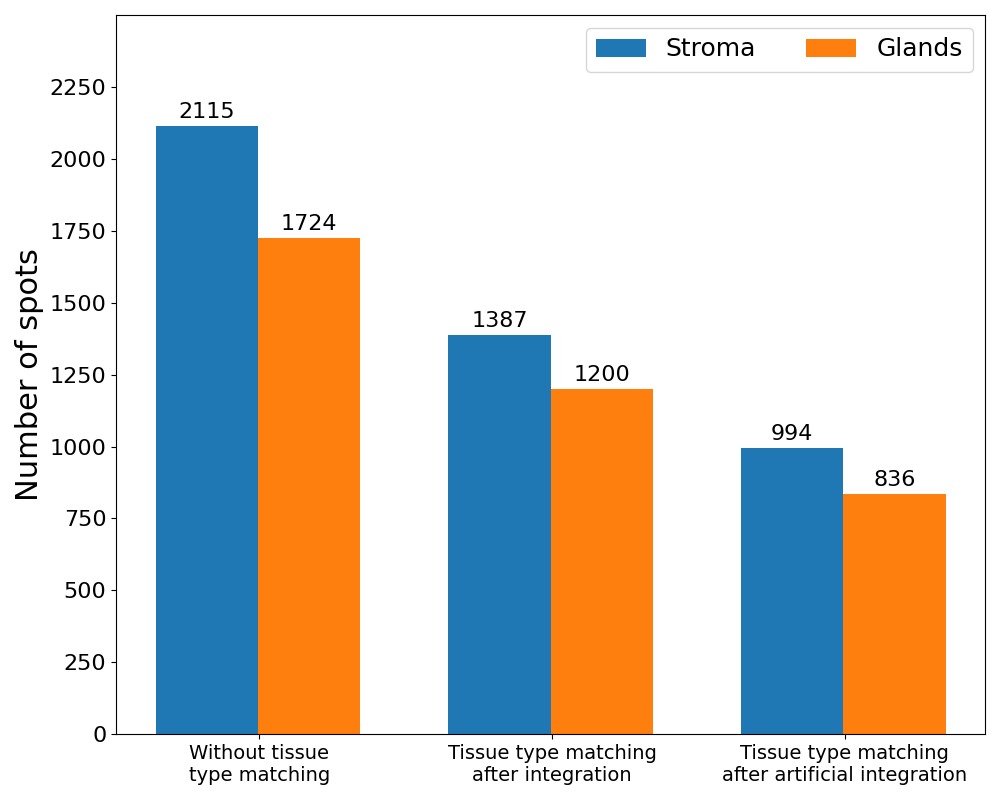

**Supplementary Figure 2.** Number of spots classified as glands or stroma with and without tissue type matching as well as after normal and artificial integration.

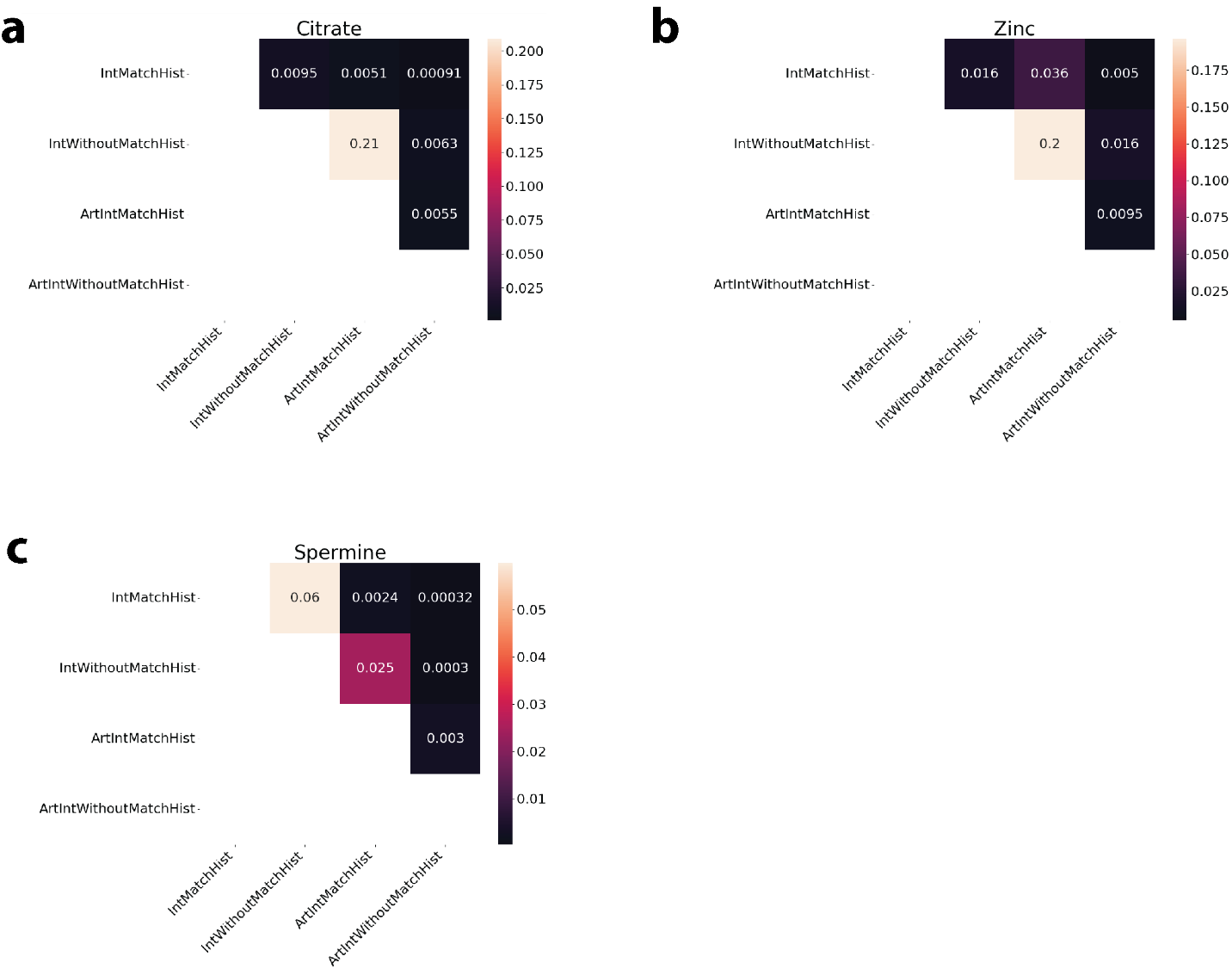
**Supplementary Figure 3.** a,b,c shows p-values for testing whether the difference of spearman correlation between two datasets is significant.

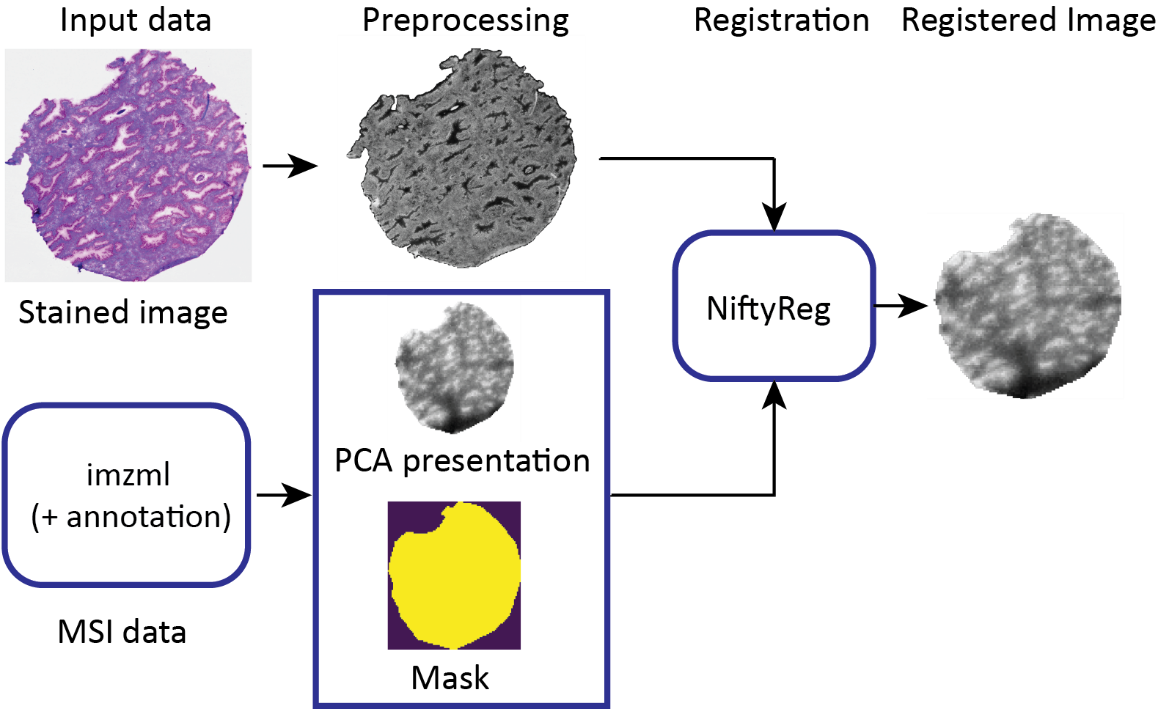

**Supplementary Figure 4.** Stained image is segmentation from background region. For the imzml file a feature image is derived based on the PCA decomposition. Then both images are padded symmetrically to a uniform shape and registered using NiftyReg. The resulting transformation is applied on the imzml data.

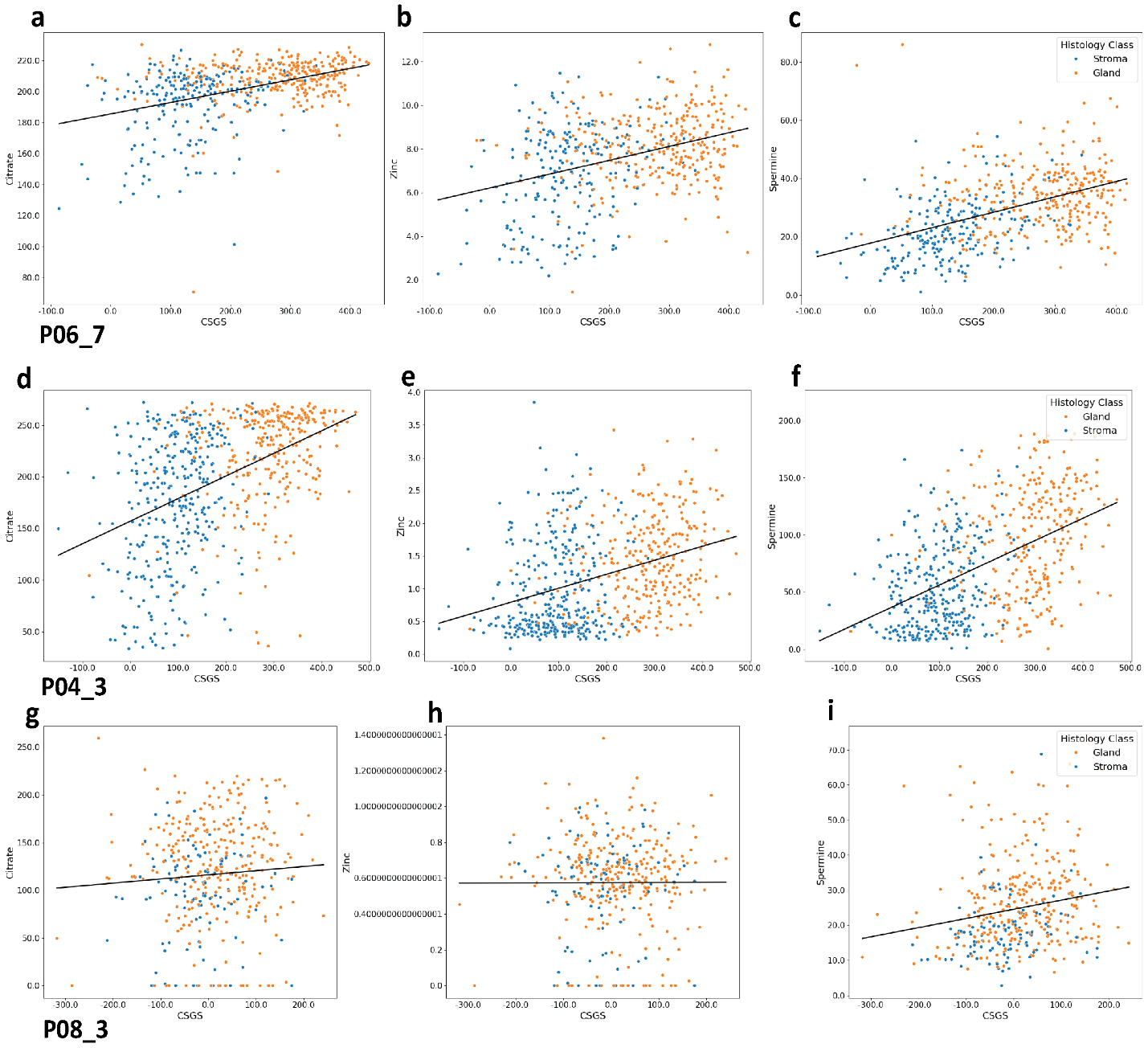

**Supplementary Figure 5.** Citrate (a, f, g), zinc (b, e, h), and spermine (c, f, i) levels plotted against CSGS score for samples P06_07 (a, b, c), P04_3 (d, e, f) and P08_3 (g, h, i) for integrated spots colored according to tissue type. Linear regression lines are shown.

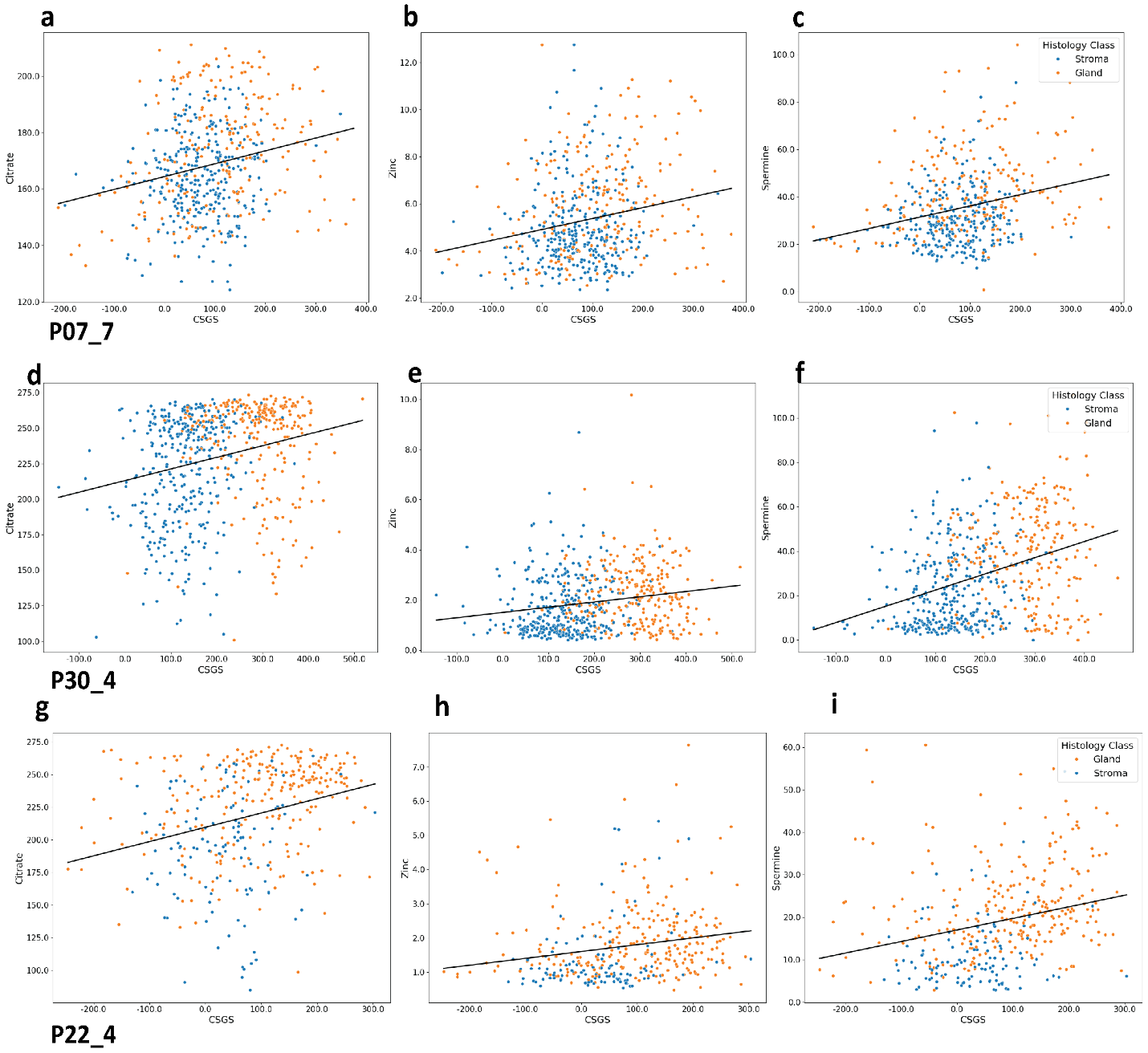

**Supplementary Figure 6.** Citrate (a, f, g), zinc (b, e, h), and spermine (c, f, i) levels plotted against CSGS score for samples P07_07 (a, b, c), P30_4 (d, e, f) and P22_4 (g, h, i) for integrated spots colored according to tissue type. Linear regression lines are shown.

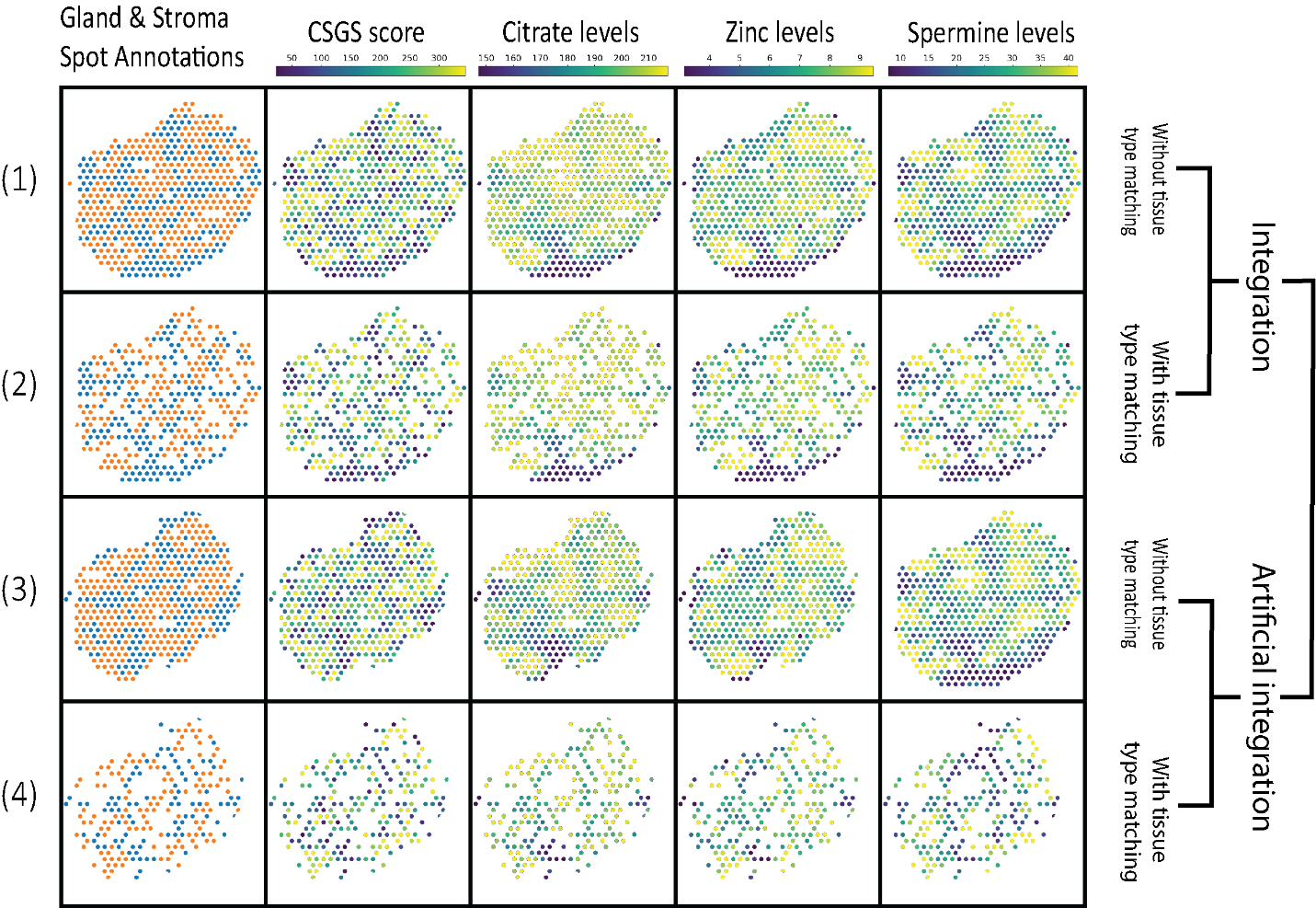

**Supplementary Figure 7.** Spot-wise distribution of gland and stroma, CSGS, citrate, zinc, and spermine for four different datasets (sample P06_7).

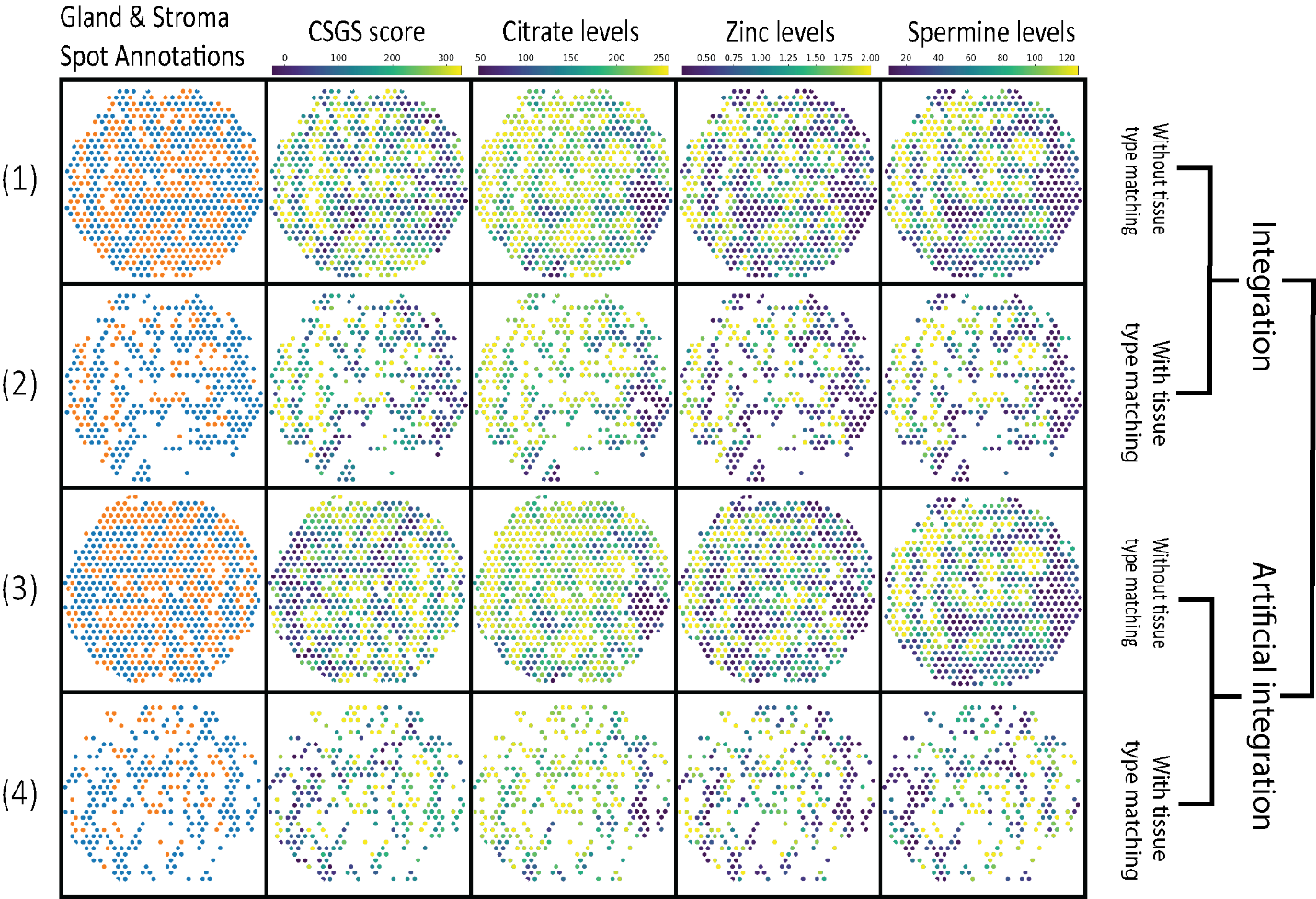
**Supplementary Figure 8.** Spot-wise distribution of gland and stroma, CSGS, citrate, zinc, and spermine for four different datasets (sample P04_3).

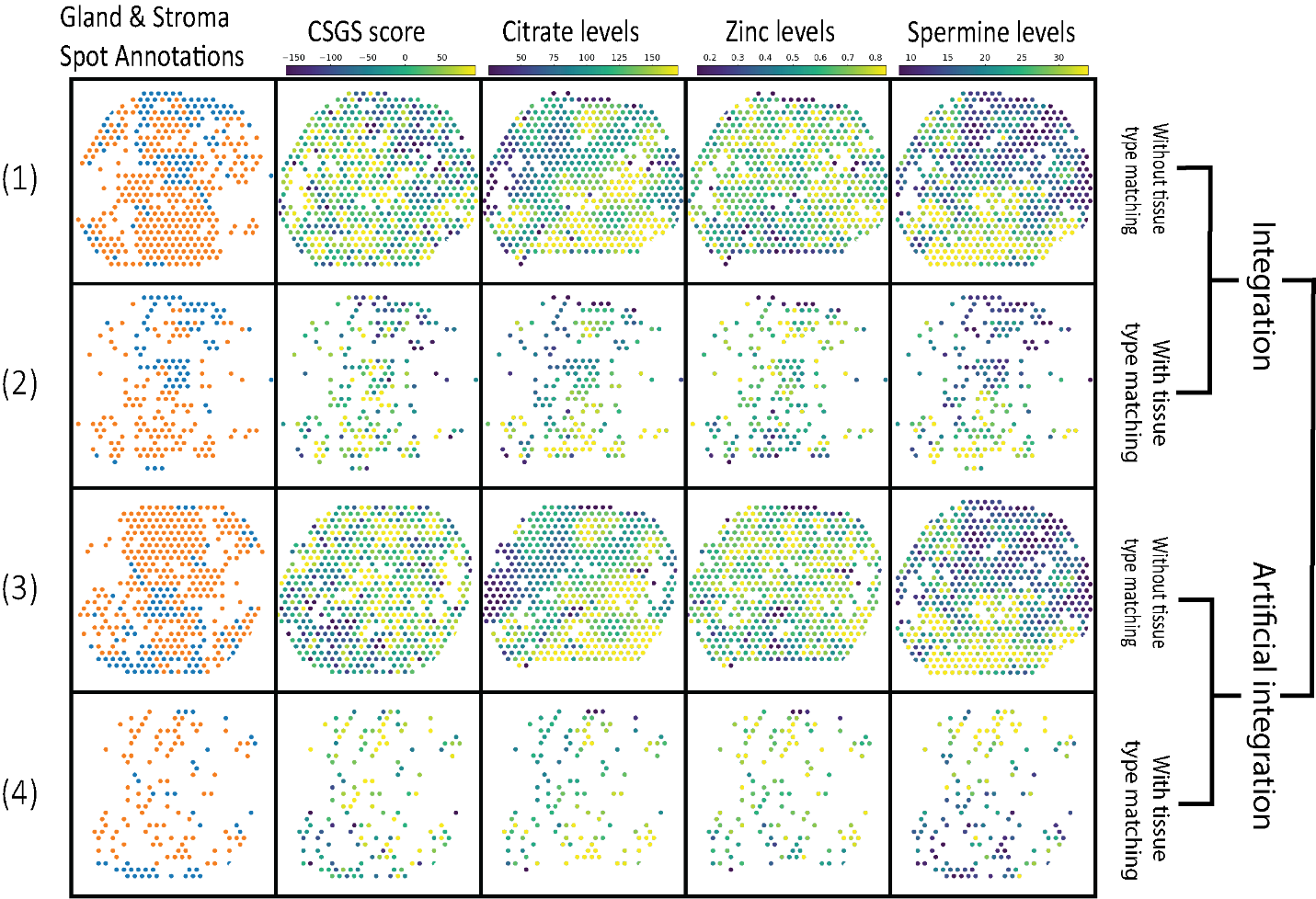
 **Supplementary Figure 9.** Spot-wise distribution of gland and stroma, CSGS, citrate, zinc, and spermine for four different datasets (sample P08_3).

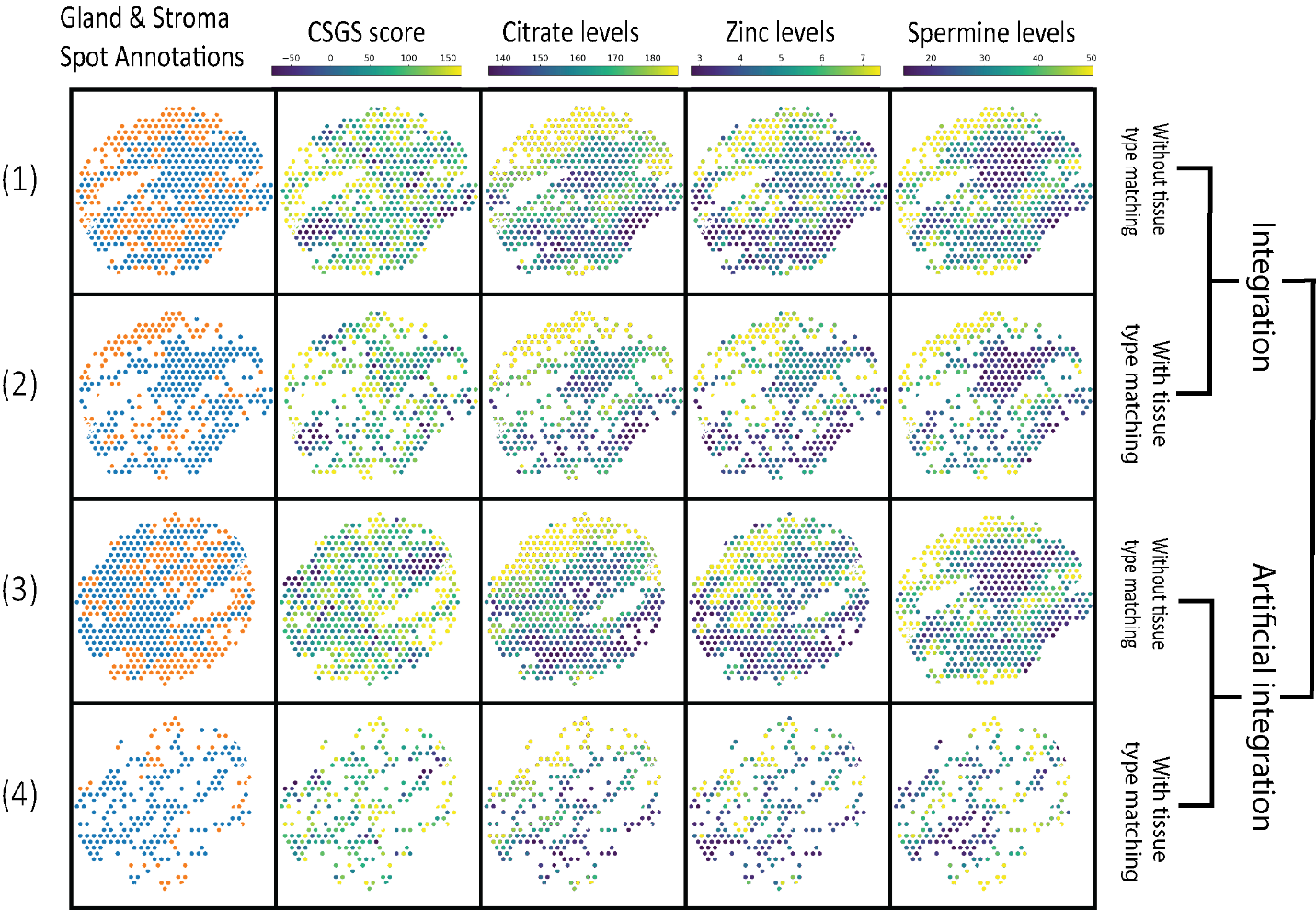
 **Supplementary Figure 10.** Spot-wise distribution of gland and stroma, CSGS, citrate, zinc, and spermine for four different datasets (sample P07_7).

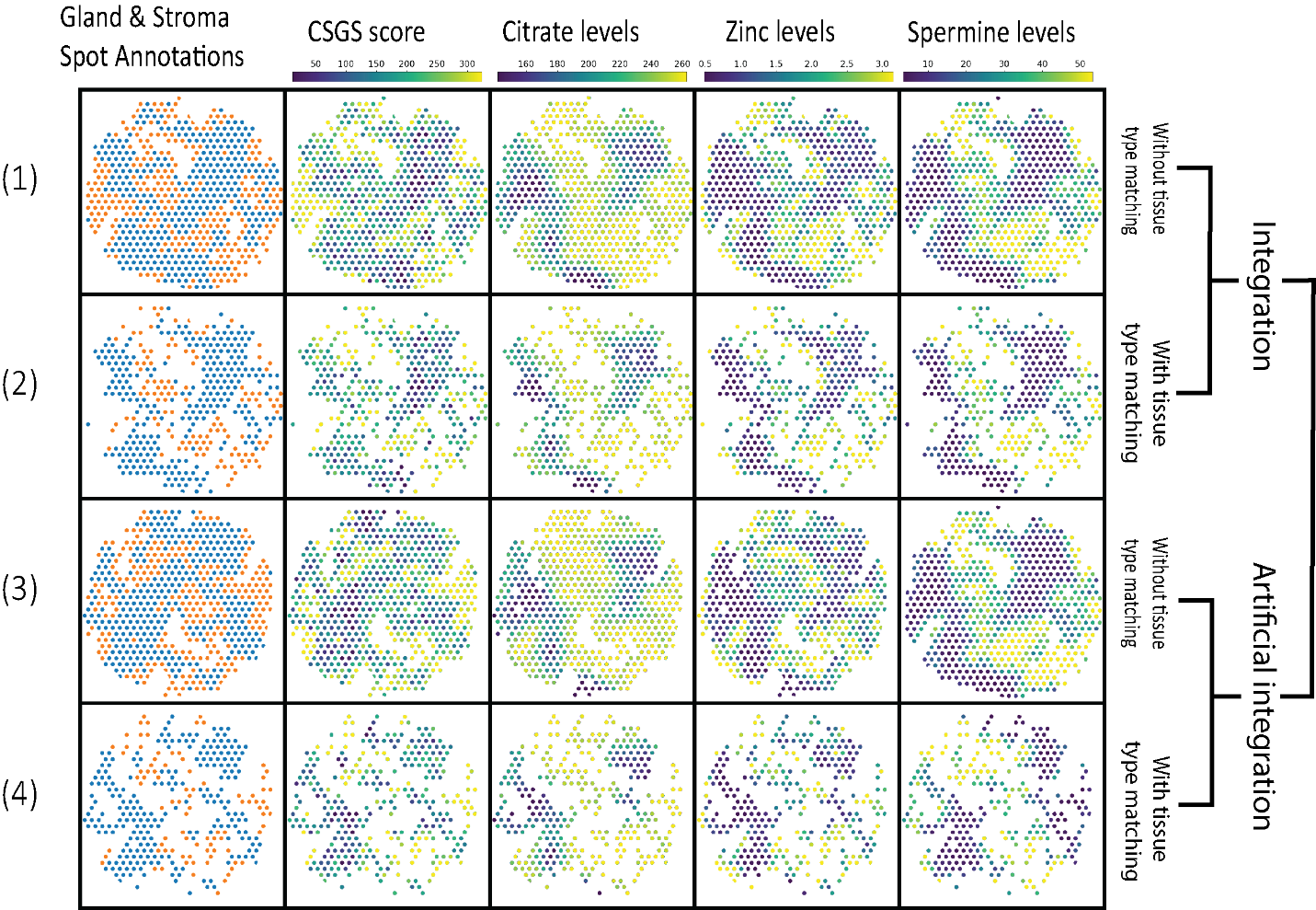

**Supplementary Figure 11.** Spot-wise distribution of gland and stroma, CSGS, citrate, zinc, and spermine for four different datasets (sample P30_4).

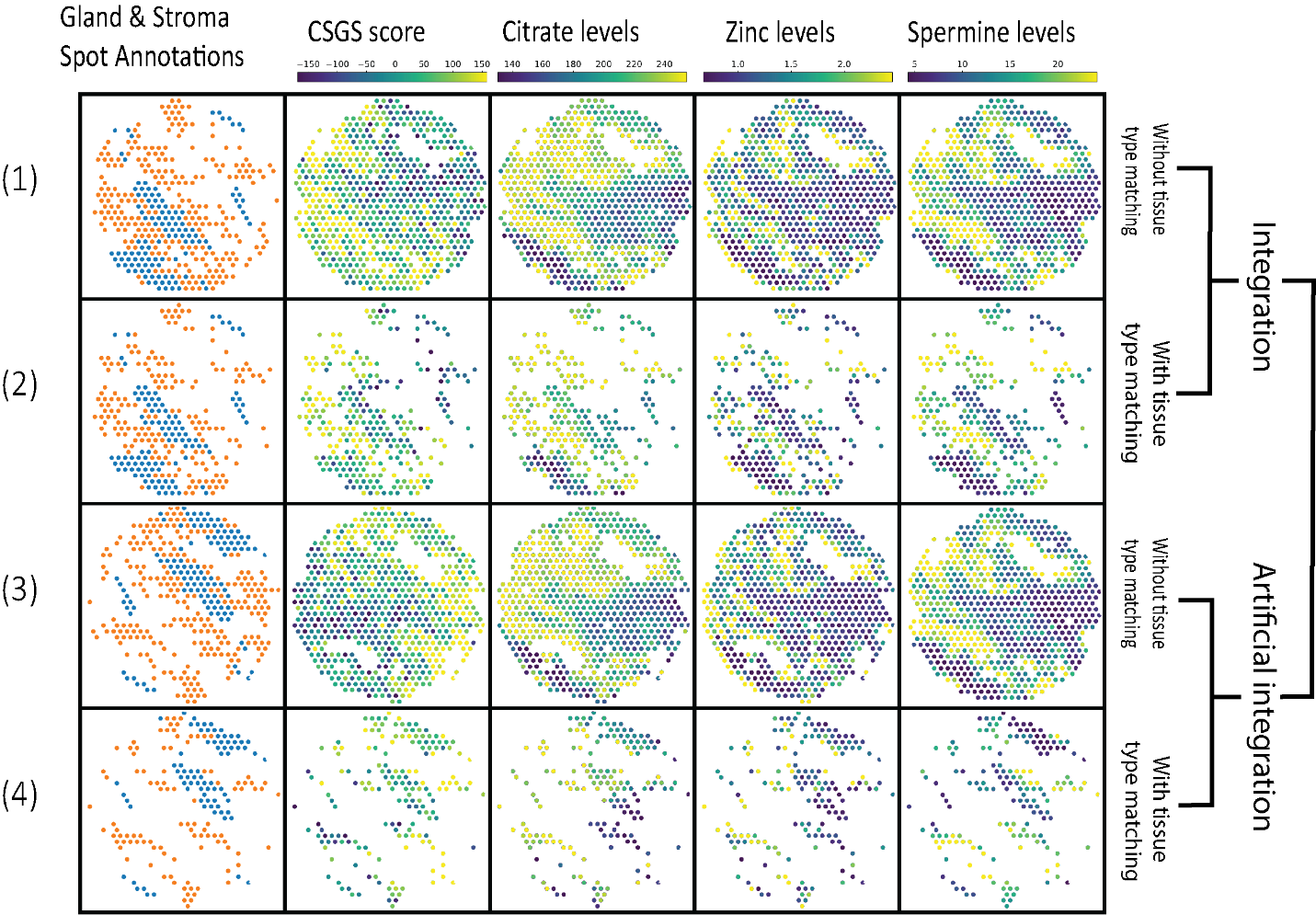
 **Supplementary Figure 12.** Spot-wise distribution of gland and stroma, CSGS, citrate, zinc, and spermine for four different datasets (sample P22_4).

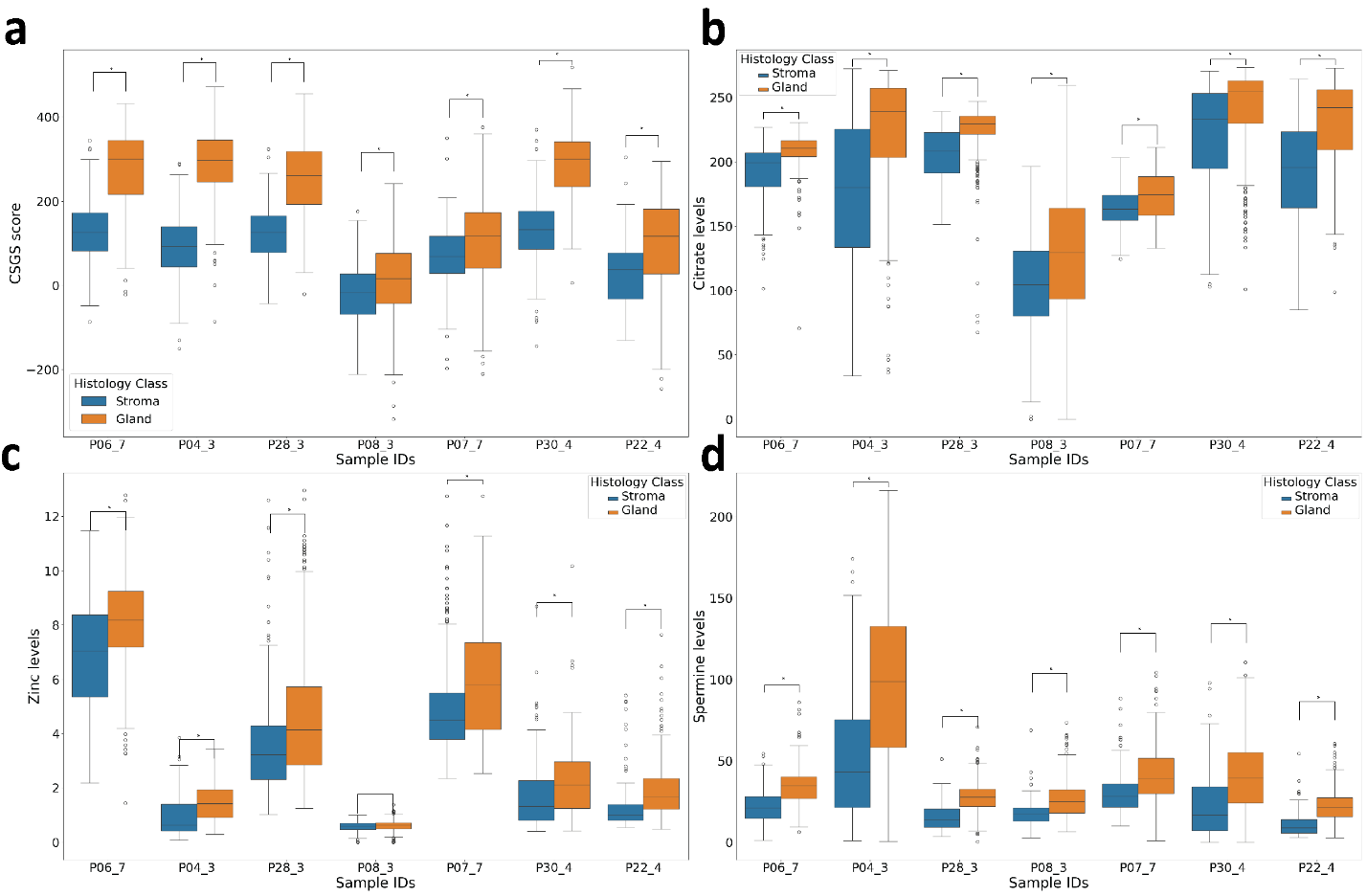

Supplementary Figure 13. Sample-wise gene scores and metabolite levels in stroma and gland spots for a) GSCS, b) citrate, c) zinc, and d) spermine.
